# ElegansBot: Development of equation of motion deciphering locomotion including omega turns of *Caenorhabditis elegans*

**DOI:** 10.1101/2023.09.26.559644

**Authors:** Taegon Chung, Iksoo Chang, Sangyeol Kim

## Abstract

Locomotion is a fundamental behavior of *Caenorhabditis elegans* (*C. elegans*). Previous works on kinetic simulations of animals helped researchers understand the physical mechanisms of locomotion and the muscle-controlling principles of neuronal circuits as an actuator part. It has yet to be understood how *C. elegans* utilizes the frictional forces caused by the tension of its muscles to perform sequenced locomotive behaviors. Here, we present a two-dimensional rigid body chain model for the locomotion of *C. elegans* by developing Newtonian equations of motion for each body segment of *C. elegans*. Having accounted for friction-coefficients of the surrounding environment, elastic constants of *C. elegans*, and its kymogram from experiments, our kinetic model (ElegansBot) reproduced various locomotion of *C. elegans* such as, but not limited to, forward-backward-(omega turn)-forward locomotion constituting escaping behavior and delta-turn navigation. Additionally, ElegansBot precisely quantified the forces acting on each body segment of *C. elegans* to allow investigation of the force distribution. This model will facilitate our understanding of the detailed mechanism of various locomotive behaviors at any given friction-coefficients of the surrounding environment. Furthermore, as the model ensures the performance of realistic behavior, it can be used to research actuator-controller interaction between muscles and neuronal circuits.

## Introduction

With only a few hundred neurons, *Caenorhabditis Elegans*(*C. elegans*) performs various behaviors such as locomotion, sleeping, reproduction, and hunting (Hall & Altun, 2008). The connectome structure among 302 neurons and 165 somatic cells of *C. elegans* was discovered by pioneering works (Cook et al., 2019; White et al., 1986). *C. elegans* is a cost-efficient and widely-used model in neuronal research. Its small body size and minimal nutritional requirements contribute to its cost efficiency. The organism matures in a shorter period, about three days, compared to other model animals such as fruit flies or mice. Its transparent body allows for easy microscopic observation of its internal structures or artificially expressed green fluorescent proteins. Moreover, due to the hermaphroditic nature of *C. elegans*, offspring mostly share the same genotype as the parent, which simplifies the multiplication of the worm population for research purposes (Hall & Altun, 2008).

*C. elegans* bends its body with a sinusoidal wave pattern when moving forward or backward. The driving force for this movement comes from the difference between perpendicular and parallel frictional forces, which it experienced from a surrounding environment. This thrust force pushes the worm along the ground surface with which the worm contacts(Boyle, 2009; Boyle et al., 2012; Hu et al., 2009; Niebur & Erdös, 1991). Even if a worm has a sinusoidal modulation generated inside it, it has difficulties in forward and backward locomotion if it does not feel the difference in frictional forces from its surroundings.

Mechanical simulators of rod-shaped animals such as *C. elegans* (Boyle et al., 2012; Niebur & Erdös, 1991), fish (Ekeberg, 1993), and snakes (Hu et al., 2009) have been used in various studies. These simulators demonstrate how the activities of muscle cells are represented as behavioral phenotypes, which are determined by signals from a neuronal circuit simulator (Boyle et al., 2012; Ekeberg, 1993; Niebur & Erdös, 1991). They also show how muscle cells return proprioceptive signals back to the neuronal circuit simulator and how animals intentionally distribute body weight for locomotion patterns (Hu et al., 2009). Similarly, the kinematic simulator of fish(Ekeberg, 1993), which has a locomotion pattern in that the animal mostly undulates in a particular direction, was used with a neuronal network simulator to model the undulation of swimming behavior. This combination of kinematic simulator and neuronal network simulator was also used to model how the locomotion pattern changes due to a surrounding environment(Boyle et al., 2012) and how the central pattern generator arises from a few cells(Boyle et al., 2012; Izquierdo & Beer, 2018).

Even though there were studies on kinematic simulation of rod-shaped animals (Boyle et al., 2012; Ekeberg, 1993; Hu et al., 2009), to our best knowledge, there was no kinetic model that reproduces complex locomotion behavior of *C. elegans*, which includes all of the various modes of locomotion of *C. elegans* such as forward locomotion, backward locomotion, and turn from experimental observations. Instead, muscle cell activities from Ansatz (Hu et al., 2009), a hypothesis of the solution, or signals from a neuronal circuit simulator (Boyle et al., 2012; Ekeberg, 1993) were applied to the kinematic simulators. A simulator should have an operational structure that imitates physical quantities from an experiment to reproduce the motion of *C. elegans* of the experiment. However, until now, no kinetic simulation has such a structure. If there is a simulator that reproduces the motion of individual experiments, analysis of the kinetics of motion of specific experiments, which provides information on the individual force that exerts on each body part of the animal, will be enabled. Also, as the kinetic simulation reproduces the motion of *C. elegans*, the behavioral phenotype that emerged from the muscle activity of neuronal circuit simulation will be more credible.

We built a Newtonian-mechanics 2-dimensional rigid body chain model of *C. elegans* to reproduce its locomotion. We incorporated its body angle, related to the contraction of the body wall muscle of *C. elegans*, into the primary operating principle of our kinetic model so that the model simulates measurable physical quantities of *C. elegans* from its experimental video. The model includes a chain of multiple rod rigid bodies, a damped torsional spring between the rigid bodies, and a control angle, which is the dynamic baseline angle from the value of the kymogram of a physical experiment. We formulated Newtonian equations of translational and rotational motion of the rigid body model and computed the numerical solution of the equations by numerical integration using semi-implicit Euler method. As a result, we were able to demonstrate trajectories and kinetics of the general locomotion of *C. elegans*, such as crawling, swimming (Vidal-Gadea et al., 2011), omega-turn, and delta-turn (Broekmans et al., 2016).

## Results

### Newton’s equation of motion for locomotion of *Caenorhabditis elegans:* How does ElegansBot work?

We introduce the simple chain model of *C. elegans*’ body. *C. elegans* has an elongated body along the head-to-tail axis. Thus, the worm’s body can be approximated as a midline extended along the anterial-posterial axis in the xy-coordinate plane (Fig. 1A). Let M (=2 μg, details in “Worm’s mass, actuator elasticity coefficient, and damping coefficient” of Supplementary Information) be the mass and L (=1 mm) be the length of the worm. Midline was approximated as n (=25) straight rods, whose ends are connected to the ends of neighboring rods (Fig. 1A). The mass, length, and moment of inertia of each rod is *m= M/n, 2r = L/n*, and *I=mr*^2^*/*3 respectively. When numbering the rods in order, with the rod at the end of the head being labeled as “1-rod” and the rod at the end of the tail being labeled as “n-rod”, let us designate the i-th rod as “i-rod”. The point where i-rod and (i+1)-rod meets is “i-joint”.

**Figure 1.**
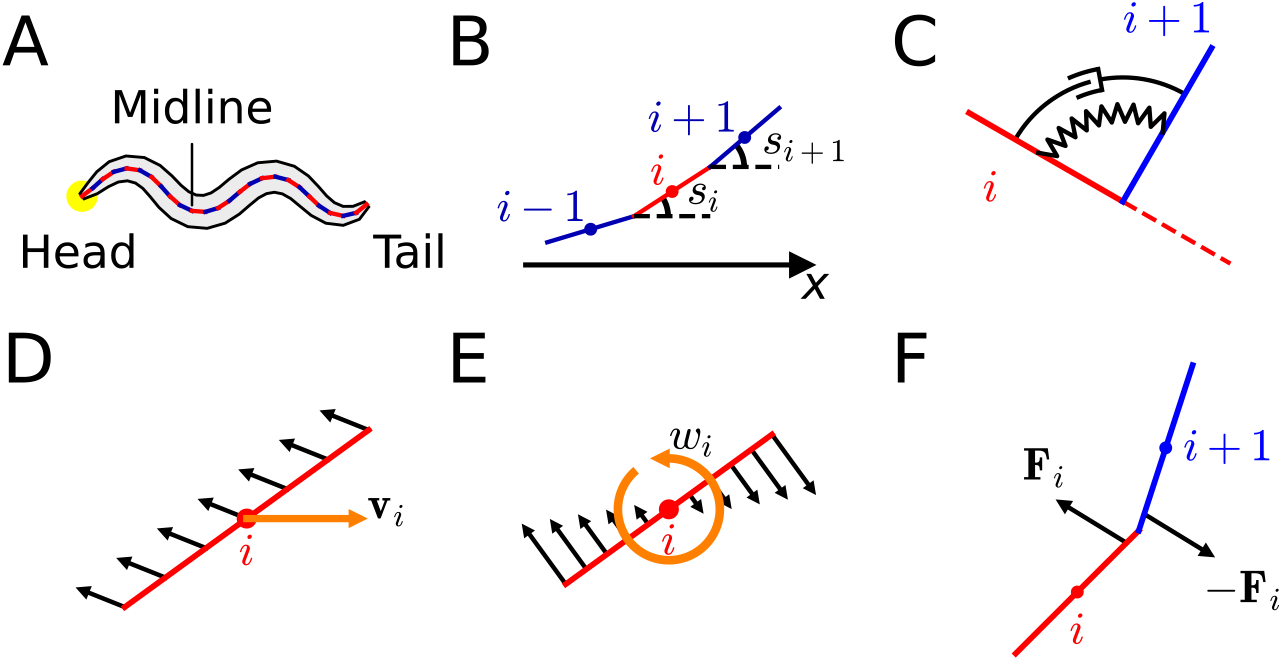
Components of ElegansBot. (A) Chain model for *C. elegans* body. (B) Rods in chain model. (C) i-actuator, which is a damped torsional spring. (D) Frictional force (black arrow) due to the translation motion of a rod. (E) Frictional force (black arrow) due to the rotational motion of a rod. (F) Joint force (black arrows) acting on i-rod and (i+1)-rod.

The motion of the worm corresponds to the motion of all the rods. To describe the motion of each rod (i-rod), we need to determine the displacement vector (*d*), velocity vector (*V*_*i*_), the angle measured counterclockwise from the positive x-axis to the tangential direction of the rod (*s*_*i*_) (Fig. 1B), and angular velocity (*ω*_*i*_) of i-rod at a given time t. However, the minimum information required to describe the motion of all rods includes the displacement vector (**d**_**c**_) and velocity vector (**V**_**c**_) of the worm’s center of mass, *s*_*i*_ and *ω*_*i*_ for each rod (Details in “Minimum information required to describe the motion of each rod” of Supplementary Information, Fig. S1).

Value of time-dependent variable such as **d**_**c**_, **V**_**c**_, *s*_*i*_, and *ω*_*i*_ at given time, t will be expressed as *\**^*(t)*^. When initial values, 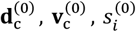, and 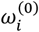 are given, the Newtonian equation of motion for acceleration, **a**_c_ and angular acceleration, {*α*_*i*_}_*i*∈{1,⋯,*n*}_ must be acquired and numerically integrated twice to find 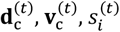, and 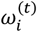 at a given time, t. To obtain the Newtonian equations of motion, we must find every force and torque acting on each rod. There are frictional force, muscle force, and joint force among types of forces acting on the rod, and there are frictional torque, muscle torque, and joint torque among types of torques whose descriptions are as follows.

The only external force acting on the worm is frictional force from a ground surface such as an agar plate or water. The frictional force is an anisotropic Stokes frictional force, with a magnitude proportional to the speed and assumed different friction coefficients in perpendicular and parallel directions (Boyle et al., 2012), which guarantees that linearity in velocity is preserved in frictional force as well (Details in “Preservation of linearity in friction” of Supplementary Information). Because of this preservation of linearity, the frictional forces of translational motion (Fig. 1D) and rotational motion (Fig. 1E) can be calculated separately and added together to find total frictional force and torque. Previously known values of the friction coefficients in perpendicular and parallel directions are used (Boyle et al., 2012).

Let the perpendicular and parallel friction coefficients be *b*_⊥_ and *b*_∥_ for a straightened worm, respectively. Each rod experiences 1/n of the frictional force the worm gets. Thus, the perpendicular and parallel friction coefficients of each rod are *b*_*⊥*_*/n, b*_∥_*/n*, respectively. The ratio of perpendicular friction coefficient to parallel friction coefficient (*b*_⊥_*/b*_∥_) is 40 in agar plate and 1.5 in water (Boyle, 2009; Boyle et al., 2012). This ratio is an important determining factor in whether the locomotion would be crawling or swimming(Boyle et al., 2012). The total frictional force that i-rod receives is 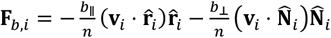 (“*⋅*”: dot product of vectors, 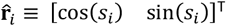 : unit vector parallel to i-rod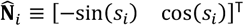 : unit vector perpendicular to i-rod), and the total frictional torque that i-rod receives is 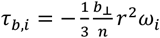 (positive or negative values are for torque pointing away from or into the paper plane, respectively.) (The proof is in “Frictional torque by rotational motion” of Supplementary Information).

Mature hermaphrodite *C. elegans* has four muscle strands at the left dorsal, right dorsal, left ventral, and right ventral part of the body, and each muscle strand has 24, 24, 23, and 24 muscle cells, respectively (White et al., 1986). Muscle cells at similar positions on the anterior-posterior axis have an activity pattern in that muscles on one side (either dorsal or ventral) cooperate, and those on the opposite side have alternative activities. (Hall & Altun, 2008).

Therefore, we modeled a group of about four muscle cells, which are left dorsal, right dorsal, left ventral, and right ventral, at the same position on the anterior-posterior axis as one actuator (Fig. 1C) so that there is a total of 24 (*≃* (24 + 24 + 23 + 24)*/*4) actuators in the worm. On i-joint of the chain, there is an actuator labeled as i-actuator. As the number of actuators is 24, we set the number of rod(n) as 25, which is one more than the number of actuators. The actuator was modeled as a damped torsional spring due to the viscoelastic characteristics of muscle(Boyle et al., 2012; Hill, 1937). If the dorsal muscles of i-actuator contract more than the ventral muscles, i-actuator will bend to the dorsal direction and vice versa. To express this phenomenon by an equation, we defined the torque that i-actuator exerts on i-rod as *τ*_*i*_ = *τ*_*κ,i*_ + *τ*_*c,i*_ that the elastic term is *τ*_*k,i*_ = *k*(*θ*_*i*_ − *θ*_ctrl,*i*_) and the damping term is *τ*_*c,i*_ = *c*(*ω*_*i*+1_ − *ω*_*i*_) where *θ*_*ctrl,i*_ is control angle, *θ*_*i*_ = *s*_*i*+1_ − *s*_*i*_, and *k* and *c* are the elasticity and damping coefficients of an actuator, respectively.

Control angle (*θ*_*ctrl,i*_) is a variable inside the elastic part of the muscle torque (*τ*_*κ,i*_), to which *τ*_*κ,i*_ drives *θ*_*i*_ close. Also, control angle (*θ*_*ctrl,i*_), which can be expressed by a heatmap (Fig. 2A, 2C, 3A, 3B), is an input value based on experimental data, a numerical model, or a neuronal network model. *τ*_*c,i*_ represents the damping effect of muscle cells and somatic cells near i-actuator. The elasticity coefficient (*κ*) and damping coefficient (*c*) of an actuator were induced from previously known values(Boyle et al., 2012) (Details in “Worm’s mass, actuator elasticity coefficient, and damping coefficient” of Supplementary Information).

**Figure 2.**
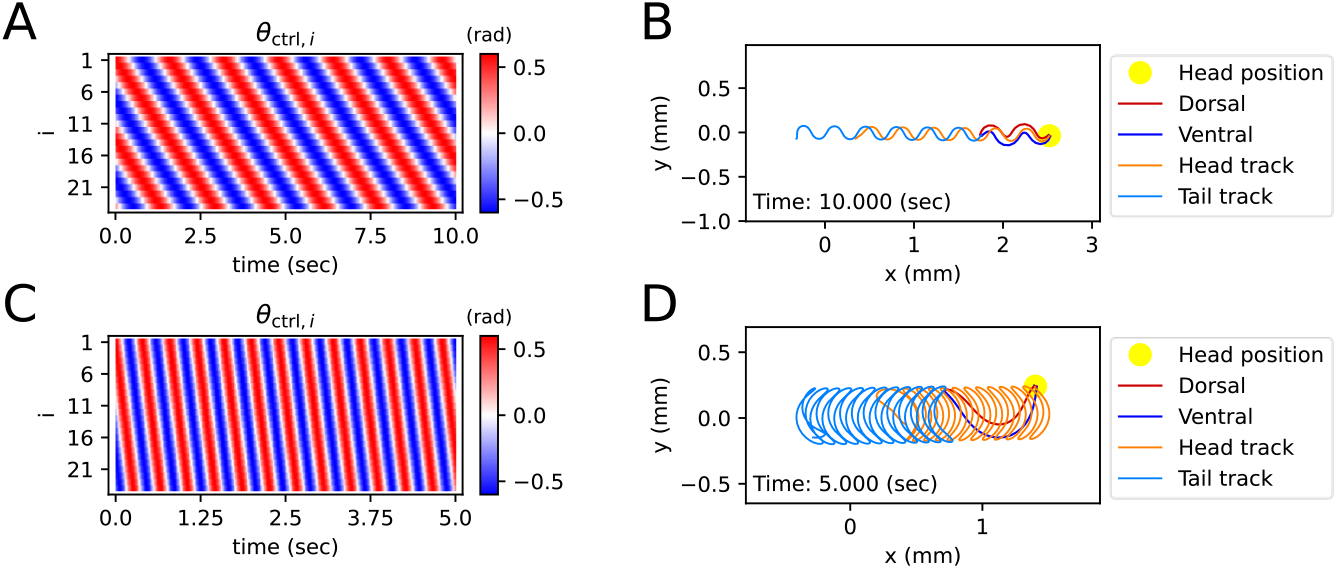
Simulated locomotion from a sine kymogram. (A) Crawling kymogram. Kymogram indicates the angle of i-joint which is located between i-rod and (i+1)-rod. Red and blue color mean i-joint bend in the dorsal and ventral directions, respectively. (B) Crawling trajectory. The yellow circle indicates the position of the worm’s head. The Orange and sky-blue lines show the worm’s head and tail trajectories, respectively. (C) Swimming kymogram. (D) Swimming trajectory

By assuming that i-rod receives torque (*τ*_*i*_) from i-actuator, and (i+1)-rod receives torque (−*τ*_*i*_), we can depict the bending that arises from the differential contraction of dorsal and ventral muscles in i-actuator. The total muscle torque that i-rod receives from damped torsional springs on both ends is *τ*_*ck,i*_ = *τ*_*i*_ − *τ*_*i*−1_. The total muscle force (**F**_*cκ,i*_) that i-rod receives from both of its ends is as follows (Details in “Proof of muscle force” of Supplementary Information, Fig. S2).

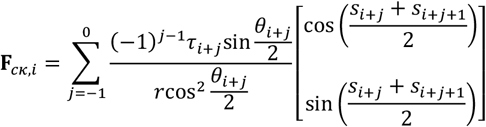

Two neighboring rods (i-rod and (i+1)-rod) are connected at i-joint. Therefore, when a force is applied to i-rod, (i+1)-rod also receives distributed force (Fig. 1F) which we name as “joint force”. The joint force that (i+1)-rod exerts on i-rod is symbolized as ***F***_*i*_ ***≡ F***_(*i*+1)*i*_ By Newton’s third law of motion about action and reaction, the joint force that i-rod exerts on (i+1)-rod is **F**_*i*(*i*+1)_ = −**F**_*i*+1)*i*_( = −**F**_*i*_ (Fig. 1F). Joint force (**F**_*i*_) can be calculated from the previously introduced given values (*s*_*i*_, **F**_*ck,i*_, **F**_*b,i*_, *τ*_*cκ,i*_, *τ*_*b,i*_) (Details in “Joint force calculation method” of Supplementary Information). When **F**_0_ = **F**_*n*_ = 0, then the total joint force that i-rod receives is **F**_joint,*i*_ = **F**_*i*_ − **F**_*i*−1_ and the total torque caused by joint force is ***τ***_joint,*i*_ = **[r**_*i*_ **×** (**F**_*i*_ + **F**_*i*+1_)] where “**×**” between two vectors means cross product.

As all forces and torques are found, 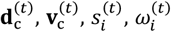 can be calculated by solving translational and rotational Newtonian equations of motion with numerical integration. The time-step (Δ*t*) used in this work is 10^−5^ sec unless otherwise noted. Because the only external force exerts on the worm is the frictional force, the equation of translational motion is 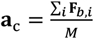. If friction coefficients (*b*_⊥_, *b*_∥_) are significantly greater than 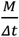, numerical integration using explicit Euler method 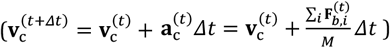 becomes unstable (Butcher, 2004). So, we tackled this instability of numerical integration by developing semi-implicit Euler Method 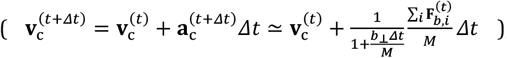, which makes numerical integration stable when any frictional coefficients greater than or equal to 0 is given (Details in “Proof of numerical integration for the translational motion of a worm using semi-implicit Euler method” of Supplementary Information).

The equation of rotational motion of i-rod is *Iα*_*i*_ = *τ*_*total,i*_ = *τ*_*b,i*_ + *τ*_*ck,i*_ + *τ*_*joint,i*_. When the friction-related value (*b*_∥_*r*^2^), elasticity-related value (*κ*Δ*t*), or damping coefficient(*c*) is significantly larger than 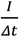, numerical integration using explicit Euler method 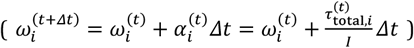 becomes unstable (Butcher, 2004). To solve this instability, we constructed a semi-implicit Euler method for rotational motion and an error-corrected equation for angular momentum (Details in “Numerical integration of the rotational motion of i-rod using semi-implicit Euler method” and “Correction formula for the rotational inertia of the entire worm” of Supplementary Information). By using these semi-implicit Euler methods, solutions for 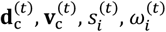 of a worm at a given time can be available for the ground surface of agar whose *b*_⊥_, *b*_∥_ are significantly larger than 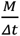, water which has smaller friction coefficients than agar, or frictionless ground surface.

### Can *C. elegans* in ElegansBot crawl or swim?

A kymogram is a heatmap that shows body angle, 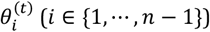 at a given time, t. By fitting a sine function to the kymogram of previous work (Vidal-Gadea et al., 2011), we obtained linear-wavenumber(after this referred to as wavenumber) and period of *C. elegans* crawling on the agar plate and swimming in water. The wavenumber (*v*) and the period (*T*) are, respectively, 1.832 and 1.6 (sec) on the agar plate and 0.667 and 0.4 (sec) in water. For both crawling and swimming, amplitude (A) was set to 0.6 (rad) arbitrarily to match the trajectory shown in the experimental video (Vidal-Gadea et al., 2011). Each kymogram of crawling (Fig. 2A) and swimming (Fig. 2C) was calculated by substituting amplitude (*A*), wavenumber (*v*), and period (*T*) into 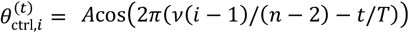.

Crawling trajectory, which performs sinusoidal locomotion in the positive x-axis direction, was obtained by inputting a crawling kymogram as 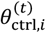 input to ElegansBot (Fig. 2B). Regarding crawling, the head track and the tail track have similar shapes. However, the tail track is more toward the negative x-axis direction than the head track. The difference between the head and tail tracks indicates that the worm pushes the ground surface by the distance between the head track and tail track to obtain thrust (Fig. 2B). Indeed, we found that the body part placed diagonally with respect to the direction of the worm’s locomotion is pushing along the ground surface (Fig. S3A). The thrust force of the worm cancels out most of the drag force, which enables the worm to move at nearly constant velocity. The average velocity of the worm is 0.208 (mm/sec), which is consistent with the known values (Cohen et al., 2012; Jung et al., 2016; Omura et al., 2012; Shen et al., 2012).

In the previous work, the worm showed swimming behavior in a water droplet on an agar plate (Vidal-Gadea et al., 2011). As the friction coefficient of water is smaller than that of agar, even though the area that the worm swept was wider during swimming than crawling, the worm did not move forward much in comparison to the area it swept (Fig. 2D). The worm gained significant momentum in the forward direction of locomotion when the body bent in the c-shape (Fig. S3B). In contrast to crawling, during swimming, the worm did not receive constant thrust force over time. Thus, the speed of the worm exhibited significant oscillations over time (Fig. S3B), and the average velocity was 0.223 (mm/sec).

### ElegansBot exhibits more complex behavior including the turn motion

Unlike previous *C. elegans* body kinematic simulation studies, our simulation can replicate the worm’s behavior using a kymogram (Fig. 3A, 3B) derived from experimental videos. We utilized open-source software, Tierpsy Tracker (Javer et al., 2018) and WormPose (Hebert et al., 2021), to obtain the kymogram input 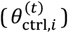 for the ElegansBot. Through simulation, we aimed to reproduce the omega-turn and delta-turn behaviors observed in the experimental videos (Broekmans et al., 2016). When we used the vertical and horizontal friction coefficients *b*_⊥_ and *b*_∥_ on agar, as proposed in the previous work (Boyle et al., 2012), the trajectory was not accurately replicated. Given that the friction coefficients could vary depending on the concentration of the agar gel, we used *b*_⊥_*/*100 and *b*_∥_*/*100 for the vertical and horizontal friction coefficients, respectively, which resulted in a better trajectory replication (Details in “Proper Selection of Friction Coefficients” in Supplementary Information, Fig. S4∼7).

**Figure 3.**
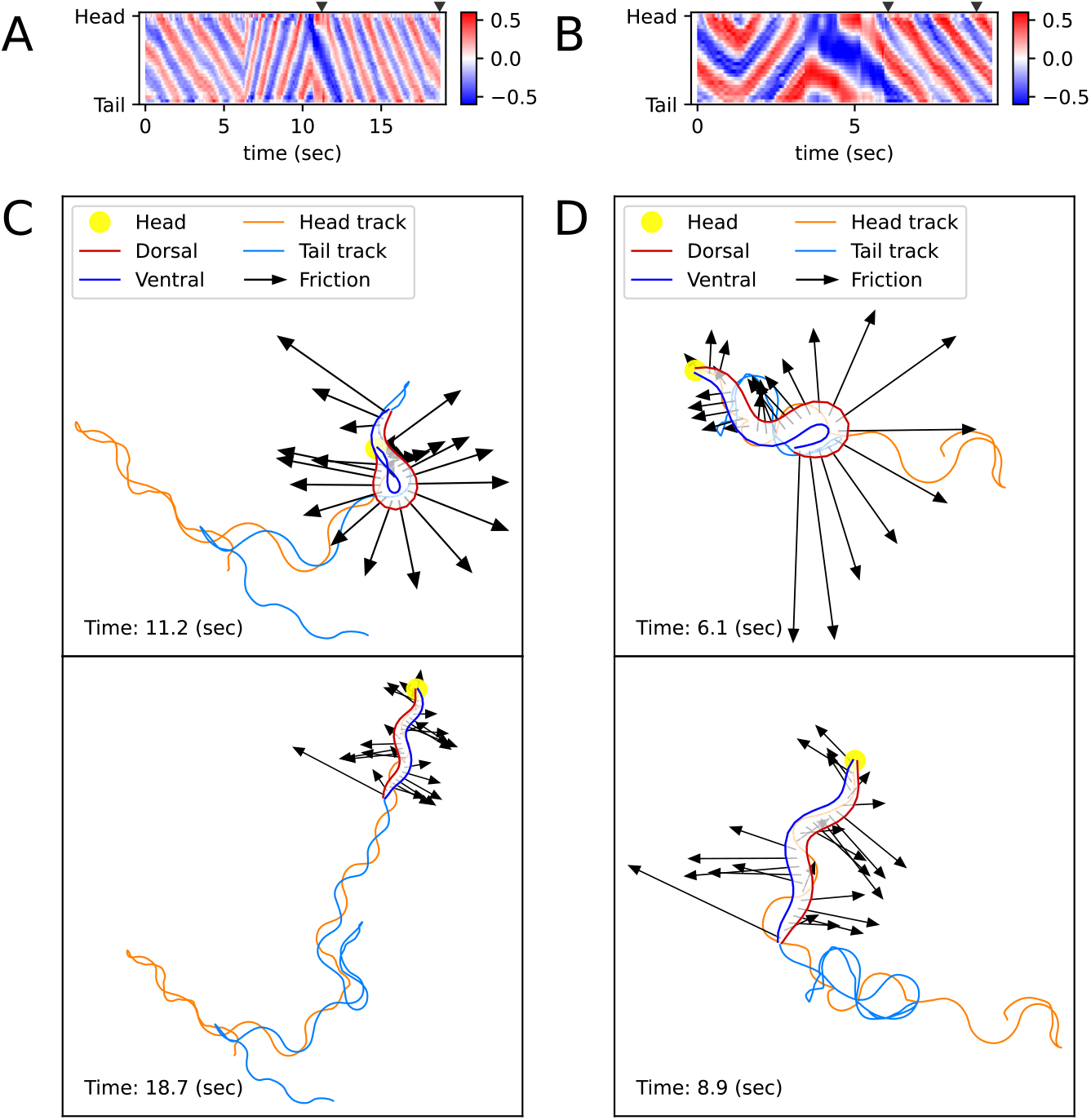
Simulated locomotion from a kymogram of a real worm locomotion video. The length and direction of a black arrow indicate the magnitude and direction of the frictional force 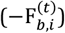 that the corresponding body part, which is the starting point of the arrow, exerts on the surface. (A) escaping behavior kymogram. Triangles over the heatmap indicate the corresponding time of snapshots shown in Figure (C). (B) Delta-turn kymogram. Triangles over the heatmap indicate the corresponding time of snapshots shown in Figure (D). (C) Escaping behavior trajectory. (D) Delta-turn trajectory. The arrow length scale is different from Figure (C) to clearly show the arrows’ directions and head and tail tracks. (Those videos are in Supplementary Information)

The trajectory (Fig. 3C, 3D, Video S1, S2) obtained from ElegansBot accurately reproduces the experimental video (Broekmans et al., 2016). The changes in the direction of movement caused by turns are well replicated. Additionally, during the omega-turn or delta-turn, the body briefly performs a deep bend, and we newly discovered the mechanism that gains significant propulsion from the deep bend region to change direction using ElegansBot (Fig. 3C, 3D). Moreover, the ElegansBot accurately reproduces not only the turns but also complex behaviors like the sequence of forward-backward-turn-forward, also known as escaping behavior.

Additionally, we calculated the mechanical power of the worm as a quantitative indicator to explain its locomotion during sequenced locomotive behavior, based on behavior classification (forward, backward locomotion, or turn, as defined in Methods). During escaping behavior, the worm produced an average power of 2,094 fW in the initial forward locomotion, followed by an average of 16,437 fW (7.85 times that of the initial forward locomotion) in backward locomotion, and an average of 11,118 fW (5.31 times that of the initial forward locomotion) during turning (Fig. 4A). After turning and resuming forward locomotion, it produced an average power of 5,480 fW (2.62 times that of the initial forward locomotion). This indicates that the worm produced more power than that of initial forward locomotion to escape sudden threats. Let’s denote the average of a quantity for all given *i* as ⟨*⟩_*i*_. At the moment the worm formed a deep bend (*t*=11.2 sec), the average magnitude of frictional force of the body part forming the deep bend 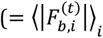 where *i*=4∼15) was 3,536 pN, compared to the average magnitude of the remaining parts 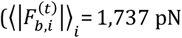 where *i*=1∼3 or *i*=16∼25), which was 2.04 times greater (Fig. 3C, Fig. 4A).

**Figure 4.**
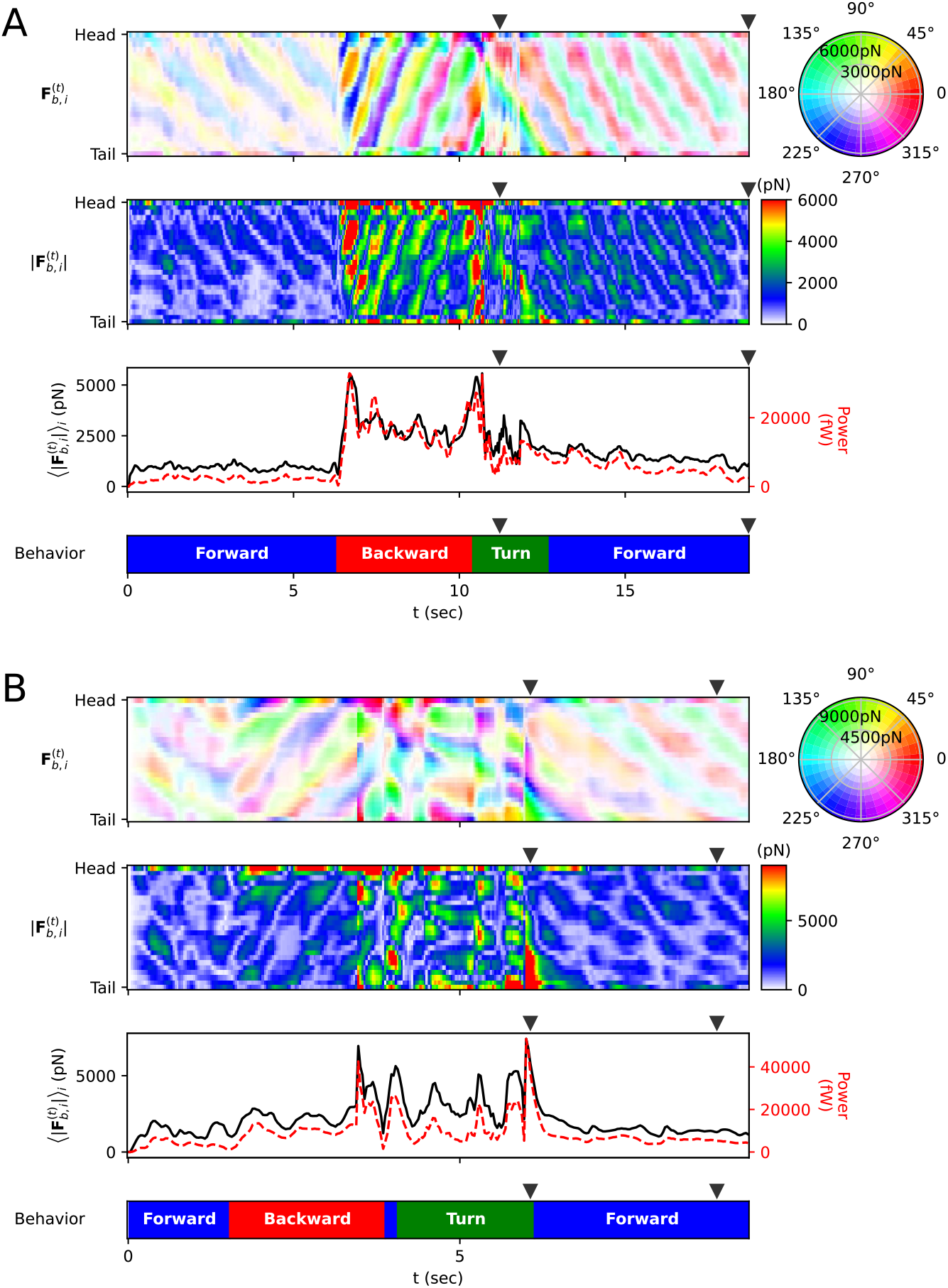
Frictional force on each rod (A) Escaping behavior. The top panel represents the frictional force 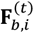 experienced by i-rod. As indicated on the color wheel to the right, the hue of this heatmap represents the direction of the force, and the saturation represents the magnitude of the force. The second panel from the top shows the magnitude of the frictional force 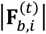. The third panel from the top represents the average 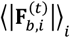 (black solid line) of each column in the middle panel and the power (red dotted line), which is the amount of energy the worm consumes per unit time. The bottom panel represents the classification of the worm’s behavior (blue: forward locomotion, red: backward locomotion, green: turn) (The definitions of behavioral categories are in Methods). The triangles over each panel indicate the corresponding time of the snapshots depicted in Fig. 3C. (B) Delta-turn. The triangles over each panel indicate the corresponding time of the snapshots depicted in Fig. 3D.

We analyzed delta-turn in the same manner. The worm produced an average power of 3,514 fW in the initial forward locomotion, followed by an average of 11,176 fW (3.18 times that of the initial forward locomotion) in subsequent backward locomotion. In the relatively short duration of forward locomotion following the backward locomotion, the worm produced average power of 17,544 fW (4.99 times that of the initial forward locomotion), and an average of 13,046 fW (3.71 times that of the initial forward locomotion) during turns (Fig. 4B). After the turn, when resuming forward locomotion, the worm produced an average power of 6,429 fW (1.83 times that of the initial forward locomotion). At the moment the worm formed a deep bend (*t*=6.1 sec), the average magnitude of frictional force of the body part 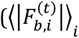where *i*=16∼25) was 10,497 pN, compared to the average magnitude of the remaining parts 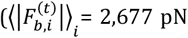 where *i*=1∼15), which was 3.92 times greater (Fig. 3D, Fig. 4B). In both escaping behavior and delta-turn, the worm consistently produced more power in the subsequent backward locomotion and turn than in the initial forward locomotion.

### ElegansBot presents body shape ensembles of *C. elegans* from a shape in water en route to agar

While there have been studies on how locomotion patterns change in agar and water by merging neural and kinematic simulations (Boyle et al., 2012), there have been none that solely used kinetic simulation to analyze how speed manifests depending on the frequency and period of locomotion. We demonstrate this aspect. We studied the locomotion speed of the worm under different friction coefficients, which represent the influence of water, agar, and intermediate frictional environment, using ElegansBot. The vertical and horizontal friction coefficients in water are *b*_water,⊥_ = 5.2 **×** 10^3^ (μg/sec) and *b*_water,∥_ = *b*_water,⊥_*/*1.5, respectively, while in agar, these values are *b*_agar,⊥_ = 1.28 **×** 10^8^ (μg/sec) and *b*_agar,∥_ = *b*_agar,⊥_/40 (Boyle et al., 2012). For environmental index σ ∈ [0,1], we have defined the vertical and horizontal friction coefficients in the environment between water (σ = 0) and agar (σ = 1) as 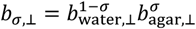 and 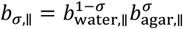, respectively.

Under an environmental index σ, for various pairs of frequency-period (*v, T*) when the control angle is 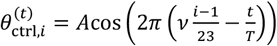 (with *A* = 0.6 (rad)), we have found the (*v, T*) that maximizes the worm’s average velocity(optimal (*v, T*)) (Fig. 5A). The optimal (*v, T*) exhibits nearly linear distribution (Fig. 5B). We noticed a transition from swimming body shape to crawling body shape as σ varies (Fig. 5, S8). The optimal (*v, T*) for σ=0(water) is (0.65, 0.4 sec), matching the actual (*v, T*) value of swimming behavior (Vidal-Gadea et al., 2011). The optimal (*v, T*) for σ=1(agar) is (1.9, 0.8 sec), and the optimal *v* (1.9) matches the actual *v* value (1.832) for crawling behavior (Vidal-Gadea et al., 2011), with the optimal T (0.8 sec) being half the actual T value (1.6 sec).

**Figure 5.**
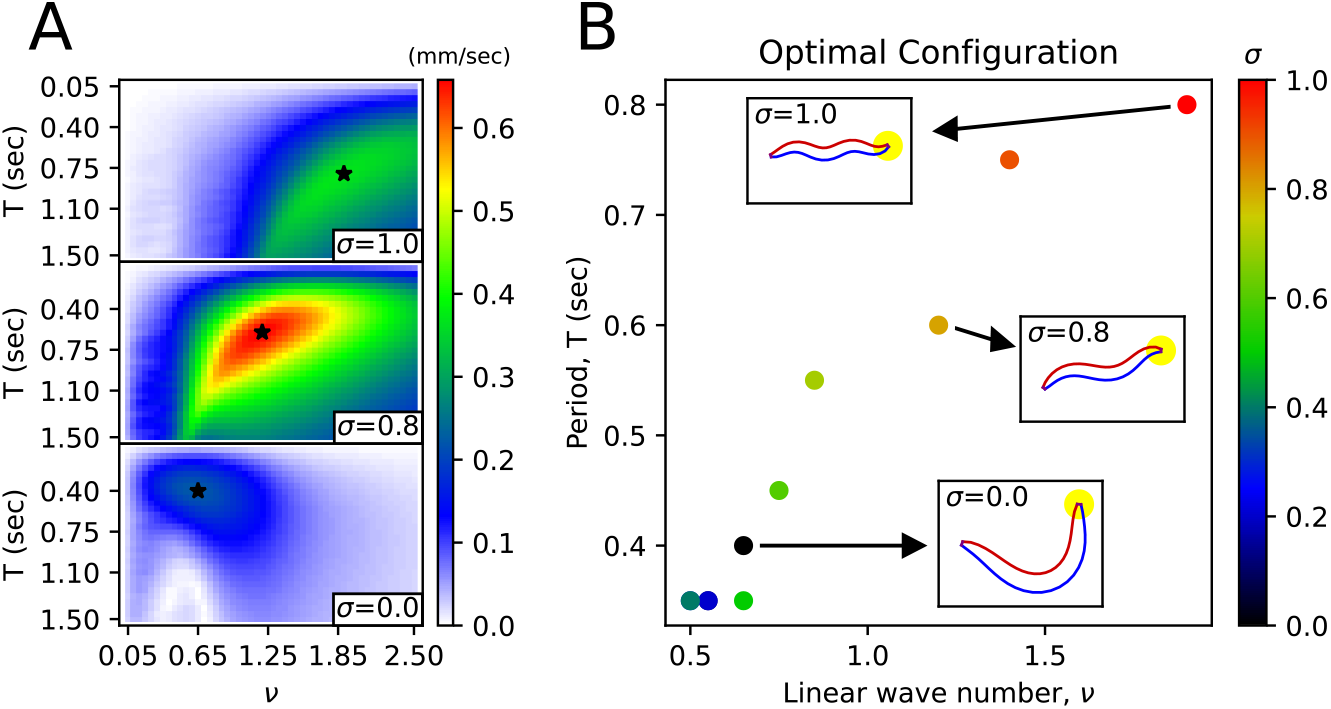
Body shape transition from the shape in water to the shape in agar. Average velocity of the worm as a function of wavenumber (*v*) and period (*T*) for a given friction coefficient. The star symbol indicates the pair of (*v, T*) that maximizes the worm’s average velocity. (B) For each environmental index σ, the pair of (*v, T*) that maximizes the worm’s average velocity. The worm figures inside the small rectangles pointed to by the arrows represent the body shape corresponding to the respective (*v, T*) pair.

We wanted to understand the impact of the environmental index σ not only on forward locomotion but also on sequenced locomotive behavior. First, we analyzed the effect of the environmental index σ on escaping behavior as follows. Let’s denote the set of a quantity for all pair of index *i* and time *t* as {*\**}_*i,t*_. When the escaping behavior kymogram input 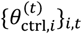 was same as Fig. 3A, we explored the effect of vertical and horizontal friction coefficients on the worm’s motion. Where σ ranged from 1.0 to 0, the trajectory varied with *σ* (Fig. S9A), and 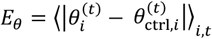 decreased as *σ* decreased (Fig. S9B). From *σ* = 1.0 to 0.1, the total absolute angular change 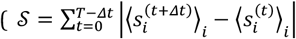 where *T* is total time of the experimental video.) increased as *σ* decreased. However, from *σ* = 0.7 to 0, *𝒮* remained constant within the error of 0.33 rad, and the total traveled distance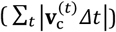 decreased as *σ* decreased. The maximum total traveled distance was at *σ* =0.8. Using the same analysis method with the kymogram input 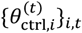 same as Fig. 3B, we analyzed the impact of the environmental index *σ* on delta-turn. Where *σ* ranged from 1.0 to 0, the trajectory varied with *σ* (Fig. S10A), and *E*_*θ*_ also decreased as *σ* decreased (Fig. S10B). From *σ* = 1 to 0.6, *𝒮* increased as *σ* decreased. From *σ* = 0.6 to 0, *𝒮* decreased as *σ* decreased. The maximum total traveled distance was at *σ* = 0.9.From *σ* = 0.9 to 0, the total traveled distance decreased as *σ* decreased.

## Discussion

### ElegansBot is an advanced kinetic simulator that reproduces *C. elegans*’ various locomotion

The known crawling speed range of *C. elegans* (Cohen et al., 2012; Jung et al., 2016; Omura et al., 2012; Shen et al., 2012) matches the speed in our simulation. The force dispersion pattern of the forward movement of a snake (Hu et al., 2009) is similar to the force dispersion pattern of crawling in our model, where the body part placed diagonally to the direction of movement generates thrust. The head and tail tracks of our simulation resemble the trace left on the agar plate by *C. elegans* during locomotion (Yeon et al., 2018), providing evidence of the mechanism where *C. elegans* moves forward by pushing along the ground surface. Given that friction and elasticity coefficients can vary between experiments, the appropriate selection of these values allows the trajectories of omega-turns and delta-turns in our simulations to match the experimental videos (Broekmans et al., 2016). Previous work (Boyle, 2009; Boyle et al., 2012) eliminated inertia from the equations of motion, but our simulation includes it, allowing calculation even in cases where inertia is significant due to low friction coefficients. Using the crawling and swimming wavenumbers and periods from the experiments (Vidal-Gadea et al., 2011), we computed sine functions to create trajectories for crawling and swimming. We also analyzed how friction forces act on the worm during crawling and swimming, studying how the worm gains propulsion. We demonstrated that we could reproduce various locomotion observed in experimental videos, such as forward-backward-(omega turn)-forward constituting escaping behavior and delta-turn navigation, by providing the kymogram obtained from representative physical values from the experimental videos, as well as the kymogram obtained from a program (Hebert et al., 2021; Javer et al., 2018) extracting the body angles from actual experimental videos into ElegansBot. Our established Newtonian equations of motion are accurate and robust, suggesting that not only does our simulation replicate the experimental videos, but it also provides credible estimates for detailed forces.

### ElegansBot will serve as a strong bridge for enhancing the knowledge in “from-synapse-to-behavior” research

Our method could be used for kinetic analysis of behaviors not covered in this paper. It could also be used when analyzing behavior changes caused by mutation or ablation experiments. Given that our simulation allows for kinetic analysis, it could be used to calculate the energy expended by the worm during locomotion, serving as an activity index. Our simulation only requires the body angles as input data, so even if the video angle shifts and trajectory information is lost, the trajectory can be recovered from the kymogram. Our simulation could also be used when studying neural circuit models of *C. elegans*. It could be used to check how signals from neural network models manifest as behaviors which is a needed function from previous work (Sakamoto et al., 2021), and it could be used when studying compound models of neural circuits and bodies. For example, when creating models that receive proprioception input based on body shape (Boyle et al., 2012; Ekeberg, 1993; Izquierdo & Beer, 2018; Niebur & Erdös, 1991), our method could be used. Finally, our method could be used in general for the broad utility to analyze the motion of rod-shaped animals like snakes or eels and to simulate the motion of rod-shaped robots.

## Methods

### 1. Frequency and wavelength of *C. elegans* locomotion

Sine function fitting was applied to the crawling and swimming kymograms (Vidal-Gadea et al., 2011) to determine the frequency and wavelength of *C. elegans* locomotion on agar and water.

### 2. *C. elegans* locomotion videos

Videos of *C. elegans*’ escaping behavior and foraging behavior were obtained from previous work (Broekmans et al., 2016). A single representative video out of a total of one hundred escaping behavior videos was used as data in this paper. Additionally, only the delta-turning portion of the foraging behavior videos was cut out and used as data in this paper.

### 3. Obtaining kymograms from video

The following method was used to extract the kymogram from the video of *C. elegans*: The body angles and midline were extracted using Tierpsy Tracker (Javer et al., 2018) from the original video where the worm is locomoting. Tierpsy Tracker failed to extract the midline of the worm when the body parts meet or the worm is coiled. The midline information of the frames successfully predicted by Tierpsy Tracker and the original video information were used as ground truth training data for a program called WormPose (Hebert et al., 2021). WormPose trained an artificial neural network to extract the body angles and midline of a coiled worm using a generative method based on the input data. The body angles were extracted from the original video using the trained WormPose program. For the frames where Tierpsy Tracker failed to extract the body angles, it was replaced with the body angles extracted by WormPose. Nonetheless, there were frames where the body angle extraction failed. If the period of failed body angle prediction was continuously less than three frames (about 0.01 seconds), the body angles for that period was predicted using linear interpolation.

### 4. Program code and programming libraries

Equations for the chain model, friction model, muscle model, and numerical integration that constitute the ElegansBot were designed from the body angle information and kymogram that change every moment of time. Python (van Rossum, 1995) version 3.8 was used to implement the equations constituting ElegansBot as a program. NumPy (Harris et al., 2020) version 1.19 was used for numerical calculations, and Numba (Lam et al., 2015) version 0.54 was used for CPU calculation acceleration. SciPy (Virtanen et al., 2020) version 1.5 was used for curve fitting and Savitzky-Golay filter (Savitzky & Golay, 1964) to classify the worm’s behavioral categories. The Matplotlib (Hunter, 2007) library was used to represent *C. elegans*’ body pose and trajectory in figures and videos. The program code used in the research can be obtained from the open database GitHub (https://github.com/taegonchung/elegansbot) or Python software repository PyPI (https://pypi.org/project/ElegansBot/), and the web live demo can be found at GitHub Page (https://taegonchung.github.io/elegansbot/). This code calculated the ElegansBot simulation of 10 seconds of simulation time in about 10 seconds of run-time on an Intel E3-1230v5 CPU.

### 5. Physical constants of the ground surface

The friction coefficient values for the ground surface where *C. elegans* crawled and swam and the elastic and damping coefficients of *C. elegans* muscles were obtained from previous work (Boyle et al., 2012). The muscle elasticity and damping coefficients were converted into coefficients for the damped torsional spring to be used in our model (Details in “Worm’s mass, actuator elasticity coefficient, and damping coefficient” of Supplementary Information).

### 6. Defining Behavioral Categories

We determined the classification of the worm’s behavior over time as follows. Let *ξ*_*i*_ ***≡*** (*i* − 1)*/*(*n* − 2) (i.e., 0 *≤ ξ*_*i*_ *≤* 1). We defined a sine function fitting for the body angle 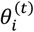 as 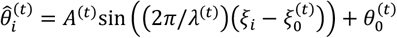 (where *A*^*(t)*^ *≥* 0 and 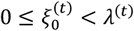). Let us denote the set of a quantity for all *i* as {*\**}_*i*_. For a given time *t*, by curve fitting the function 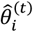 to the set of body angles 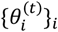, we can obtain the parameters 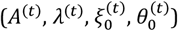 (Fig. S11). For curve fitting, we used ‘curve_fit’ from the scipy library (Virtanen et al., 2020). ‘curve_fit’ requires the function to be fitted, the data, and the initial guess values of the function parameters. Therefore, we determined the initial guess values 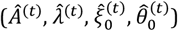 as follows. For *t* = 0, we calculated 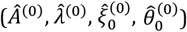 using the following equations:

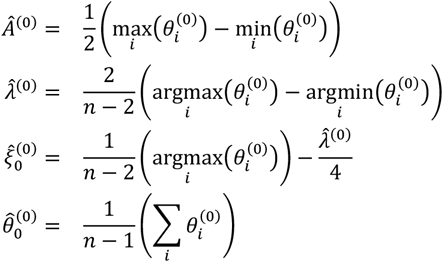

Using these initial guess values, we curve fitted 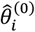 to 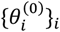 to obtain 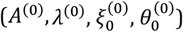. For *t ≥* Δ*t*, we obtained the initial guess values for curve fitting as 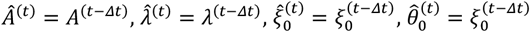. Then, using these initial guess values, we curve fitted 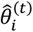 to 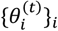 to obtain 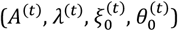. Since the phase 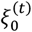 is not continuous for all time *t*, we defined a continuous value 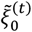 for all time *t* as follows (Fig. S11):

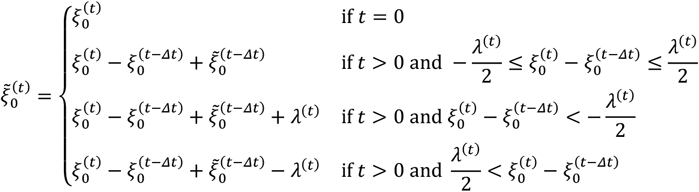

To obtain the derivatives of the noise-reduced smoothed values 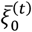 and 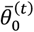 for the raw data 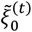 and 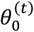, respectively, we applied a Savitzky-Golay filter (Savitzky & Golay, 1964; Virtanen et al., 2020). This filter, set with a smoothing time window of 0.5 second, a polynomial order of 2, and a derivative order of 1, yielded 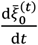 and 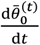. We then calculated the temporal integrals 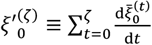 and 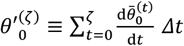. Let us denote the average of a quantity for all time t as ⟨*⟩_*t*_. We calculated 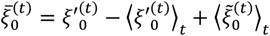 and 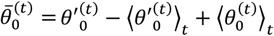 (Fig. S11). Finally, we defined the worm’s behavior classification as turn when 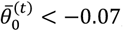 as forward locomotion when 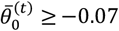 and 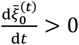, and as backward locomotion when 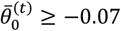 and 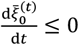 (Fig. S11).

## Acknowledgement

This work is supported by the p-CoE program of DGIST 23-CoE-BT-01. We also acknowledge Prof. Kyuhyung Kim for fruitful discussions on experiments of C. elegans.

