## Supplementary Information for "ElegansBot: Development of equation of motion deciphering locomotion including omega turns of *Caenorhabditis elegans*"

Taegon Chung, Iksoo Chang, and Sangyeol Kim
Department of Brain Sciences,
Daegu Gyeongbuk Institute of Science and Technology, Korea

2024 January

### Supplementary Information

#### Worm's mass, actuator elasticity coefficient, and damping coefficient

1. **Mass**

   The maximum radius of the worm is 40μm (Boyle et al., 2012). Assuming that the border surrounding the cross-section parallel to the anterior-posterior axis is a sine function, the average radius is $\gamma = 40 \times 2/\pi \simeq 25$μm. Assuming that the worm is a cylindrical body with a bottom surface radius of 25μm, the volume of the worm is $1\text{mm}(0.025\text{mm})^2\pi \simeq 0.002\text{mm}^3$, and the density of the worm is $\simeq 1000\mu\text{g/mm}^3$ (Reina et al., 2013), so the weight of the worm is $M = 1000\mu\text{g/mm}^3 \times 0.002\text{mm}^3 = 2\mu\text{g}$.

2. **Torque elasticity coefficient**

   In previous research, the muscle elasticity and damping coefficient were designed as functions of the input signal(Boyle et al., 2012). The maximum value of this muscle elasticity coefficient is $k_{\text{max}} = 2.8 \cdot 10^8[\mu\text{g/sec}^2]$, and the maximum value of the muscle damping coefficient is $(k_{\text{max}}/5.6) \cdot (1\text{sec})$. When $\theta_{i-1} = \theta_i = \theta_{i+1} = 0$ and the length of the moment arm where the muscle exerts force is equal to the average radius $\gamma$ of the worm, the change in the elastic torque due to the change in $\theta_i$ is the torque elasticity coefficient $\kappa$, so $\kappa = \frac{\mathrm{d}\tau_{\kappa,i}}{\mathrm{d}\theta_i} = \frac{\mathrm{d}}{\mathrm{d}\theta_i}\left(\gamma\left(k\left(2\gamma\tan(\theta_i/2)\right)\right) \simeq k\gamma^2\right) = 1.75 \cdot 10^5\mu\text{g} \cdot \text{mm}^2/(\text{sec}^2 \cdot \text{rad})$. In ElegansBot, it was assumed that this $\kappa$ value is constant regardless of $\theta_{i-1}$ , $\theta_i$ , $\theta_{i+1}$. In the same way, the torque muscle damping coefficient is $c = (1/5.6) \cdot 1.75 \cdot 10^5\mu\text{g} \cdot \text{mm}^2/(\text{sec} \cdot \text{rad})$.

#### Minimum information required to describe the motion of each rod

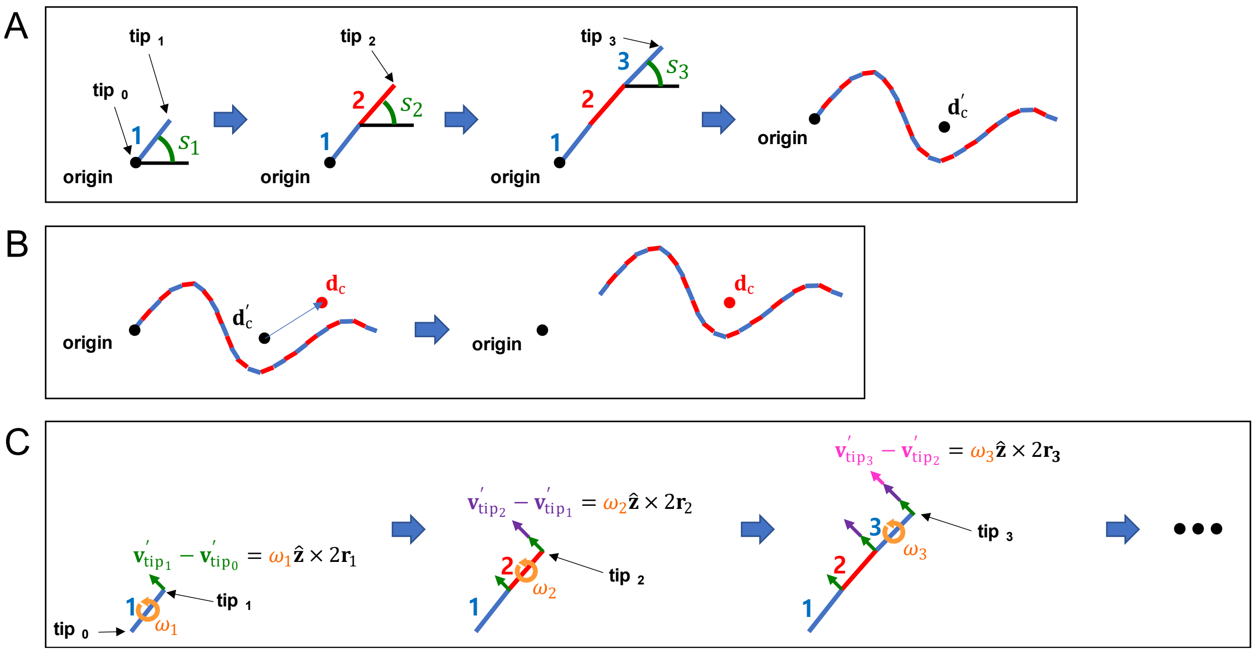


**Figure S1.** Method of compressing motion state information. (A) Method of calculating the relative position of the rod. (B) Method of calculating the absolute position of the rod. (C) Method of calculating the relative velocity of the rod.

##### 1. Information required to describe the movement of the rod

To describe the motion of all rods, it is necessary to know the position ($\mathbf{d}_i$), velocity ($\mathbf{v}_i$), angle ($s_i$) measured counterclockwise from the positive x-axis to the direction of vector $\mathbf{r}_i$, and angular velocity($\omega_i$) of every i-rod. However, knowing only the position and velocity of the worm ($\mathbf{d}_c$ , $\mathbf{v}_c$) and the angle and angular velocity of every i-rod ($s_i$ , $\omega_i$) is sufficient to describe the motion of all rods.

##### 2. Boundary conditions for the position of the rod

The center of mass of the worm is $\mathbf{d}_c = [x_c \quad y_c]^\mathsf{T} = (\sum_{i=1}^{n} m\, \mathbf{d}_i)/M = (\sum_{i=1}^{n} \mathbf{d}_i)/n$, and the vector parallel to i-rod is $\mathbf{r}_i \equiv r[\cos(s_i) \quad \sin(s_i)]^\mathsf{T} = r\hat{\mathbf{r}}_i$. If the position vector where i-rod and (i+1)-rod meet is $\mathbf{d}_{\mathrm{tip}_i}$, and the free end of 1-rod and n-rod are $\mathbf{d}_{\mathrm{tip}_0}$ and $\mathbf{d}_{\mathrm{tip}_n}$, then $\mathbf{d}_{\mathrm{tip}_i} = \mathbf{d}_i + \mathbf{r}_i = \mathbf{d}_{i+1} - \mathbf{r}_{i+1}$ (Fig. S1A).

##### 3. Method of calculating the relative and absolute position of the rod

If the relative coordinate with $\mathbf{d}_{\mathrm{tip}_0}$ as the origin is designated as $\mathbf{d}'$, then the following equations satisfy.

$$
\begin{aligned}
\mathbf{d}'_{\mathrm{tip}_0} &= \begin{bmatrix} 0 \\ 0 \end{bmatrix} \\
\mathbf{d}'_{\mathrm{tip}_i} &= \sum_{j=1}^{i} 2\,\mathbf{r}_j \\
\mathbf{d}'_i &= \frac{\mathbf{d}'_{\mathrm{tip}_i} + \mathbf{d}'_{\mathrm{tip}_{i-1}}}{2} \\
\mathbf{d}'_{\mathrm{c}} &= \frac{1}{n}\sum_{i=1}^{n} \mathbf{d}'_i \\
\mathbf{d}_i &= \mathbf{d}'_i - \mathbf{d}'_{\mathrm{c}} + \mathbf{d}_{\mathrm{c}}
\end{aligned}
$$

Thus, $\mathbf{d}_i$ can be calculated from $\mathbf{d}_{\mathrm{c}}$ and $s_i$ (Fig. S1B).

#### 4. Method of calculating the relative and absolute velocity of the rod

Based on the relationship between the worm's momentum and the rods' momentum, $\mathbf{v}_{\mathrm{c}} = (\sum_{i=1}^{n} m\,\mathbf{v}_i)/M = (\sum_{i=1}^{n} \mathbf{v}_i)/n$. In the same way as the location information compression, the relative velocity vector $\mathbf{v}'$ with $\mathbf{v}_{\mathrm{tip}_0}$ as the origin has the following relationships.

$$
\begin{aligned}
\mathbf{v}'_{\mathrm{tip}_0} &= \begin{bmatrix} 0 \\ 0 \end{bmatrix} \\
\mathbf{v}'_{\mathrm{tip}_i} &= \sum_{j=1}^{i} \omega_j\,\hat{\mathbf{z}} \times (2\mathbf{r}_j) \\
\mathbf{v}'_i &= \frac{\mathbf{v}'_{\mathrm{tip}_i} + \mathbf{v}'_{\mathrm{tip}_{i-1}}}{2} \\
\mathbf{v}'_{\mathrm{c}} &= \frac{1}{n}\sum_{i=1}^{n} \mathbf{v}'_i \\
\mathbf{v}_i &= \mathbf{v}'_i - \mathbf{v}'_{\mathrm{c}} + \mathbf{v}_{\mathrm{c}}
\end{aligned}
$$

Thus, $\mathbf{v}_i$ can be calculated from $\mathbf{v}_{\mathrm{c}}$ , $s_i$, and $\omega_i$ (Fig. S1C).

#### Preservation of linearity in friction

The velocity $\mathbf{v}$ of an arbitrary point particle which is included by i-rod can be decomposed into two velocity components $\mathbf{v} = \mathbf{v}_\alpha + \mathbf{v}_\beta$. In this case, if the object is subject to an anisotropic Stokes friction, there is linearity between the friction $\mathbf{F}_{b,\alpha}$, $\mathbf{F}_{b,\beta}$ obtained from each velocity component $\mathbf{v}_\alpha$, $\mathbf{v}_\beta$ and the friction $\mathbf{F}_b$ obtained from velocity $\mathbf{v}$.

$$\begin{aligned}
\mathbf{F}_{b,\alpha} &= -b_\parallel n^{-1}(\mathbf{v}_\alpha \cdot \hat{\mathbf{r}}_i)\hat{\mathbf{r}}_i - b_\perp n^{-\mathbf{1}}\left(\mathbf{v}_\alpha \cdot \hat{\mathbf{N}}_i\right)\hat{\mathbf{N}}_i \\
\mathbf{F}_{b,\beta} &= -b_\parallel n^{-1}\left(\mathbf{v}_\beta \cdot \hat{\mathbf{r}}_i\right)\hat{\mathbf{r}}_i - b_\perp n^{-\mathbf{1}}\left(\mathbf{v}_\beta \cdot \hat{\mathbf{N}}_i\right)\hat{\mathbf{N}}_i \\
\mathbf{F}_b &= -b_\parallel n^{-1}(\mathbf{v} \cdot \hat{\mathbf{r}}_i)\hat{\mathbf{r}}_i - b_\perp n^{-\mathbf{1}}\left(\mathbf{v} \cdot \hat{\mathbf{N}}_i\right)\hat{\mathbf{N}}_i \\
&= -b_\parallel n^{-\mathbf{1}}\left(\left(\mathbf{v}_\alpha + \mathbf{v}_\beta\right) \cdot \hat{\mathbf{r}}_i\right)\hat{\mathbf{r}}_i - b_\perp n^{-\mathbf{1}}\left(\left(\mathbf{v}_\alpha + \mathbf{v}_\beta\right) \cdot \hat{\mathbf{N}}_i\right)\hat{\mathbf{N}}_i \\
&= -b_\parallel n^{-\mathbf{1}}(\mathbf{v}_\alpha \cdot \hat{\mathbf{r}}_i)\hat{\mathbf{r}}_i - b_\perp n^{-\mathbf{1}}\left(\mathbf{v}_\alpha \cdot \hat{\mathbf{N}}_i\right)\hat{\mathbf{N}}_i \\
&\quad -b_\parallel n^{-1}\left(\mathbf{v}_\beta \cdot \hat{\mathbf{r}}_i\right)\hat{\mathbf{r}}_i - b_\perp n^{-1}\left(\mathbf{v}_\beta \cdot \hat{\mathbf{N}}_i\right)\hat{\mathbf{N}}_i \\
&= \mathbf{F}_{b,\alpha} + \mathbf{F}_{b,\beta}
\end{aligned}$$

#### Frictional torque by rotational motion

Let us denote variable $\rho$ as the distance from the center of i-rod measured along the direction of vector $\hat{\mathbf{r}}_i$. It is to be noted that $\rho$ is within the range $[-r, r]$. For an infinitesimal $\mathrm{d}\rho$ where $0 < \mathrm{d}\rho \ll 1$, the moment arm vector for the infinitesimal interval $[\rho - \mathrm{d}\rho/2, \rho + \mathrm{d}\rho/2]$ (hereafter referred to as the infinitesimal interval $\rho$) from the center of i-rod is $\rho\hat{\mathbf{r}}_i$. The coefficient of friction for the infinitesimal interval $\rho$ is:

$$\frac{b_\perp}{n}\frac{\mathrm{d}\rho}{2r}$$

The velocity component due to the rotational motion of the infinitesimal interval $\rho$ is $\rho\omega_i\hat{\mathbf{N}}_i$. Therefore, the frictional force received by the infinitesimal interval $\rho$ due to rotational motion is:

$$-\frac{b_\perp}{n}\frac{\mathrm{d}\rho}{2r}\rho\omega_i\hat{\mathbf{N}}_i = -\frac{1}{2}\frac{b_\perp}{nr}\omega_i\rho\mathrm{d}\rho\hat{\mathbf{N}}_i$$

The torque received by the infinitesimal interval $\rho$ due to rotational motion is:

$$\begin{aligned}
\mathrm{d}\boldsymbol{\tau} &= (\rho\hat{\mathbf{r}}_i) \times \left(-\frac{1}{2}\frac{b_\perp}{nr}\omega_i\rho\mathrm{d}\rho\hat{\mathbf{N}}_i\right) \\
&= -\frac{1}{2}\frac{b_\perp}{nr}\omega_i\rho^2\mathrm{d}\rho\hat{\mathbf{r}}_i \times \hat{\mathbf{N}}_i \\
&= -\frac{1}{2}\frac{b_\perp}{nr}\omega_i\rho^2\mathrm{d}\rho\hat{\mathbf{z}}
\end{aligned}$$

The total frictional torque received by i-rod due to rotational motion is

$$\begin{aligned}
\boldsymbol{\tau}_{b,i} &= \int_{-r}^{r}\mathrm{d}\boldsymbol{\tau} \\
&= \int_{-r}^{r} -\frac{1}{2}\frac{b_\perp}{nr}\omega_i\rho^2\mathrm{d}\rho\hat{\mathbf{z}} \\
&= -\frac{1}{2}\frac{b_\perp}{nr}\omega_i\left[\frac{1}{3}\rho^3\right]_{-r}^{r}\hat{\mathbf{z}} \\
&= -\frac{1}{2}\frac{b_\perp}{nr}\omega_i\frac{2}{3}r^3\hat{\mathbf{z}} \\
&= -\frac{1}{3}\frac{b_\perp}{n}r^2\omega_i\hat{\mathbf{z}}
\end{aligned}$$

#### Proof of muscle force

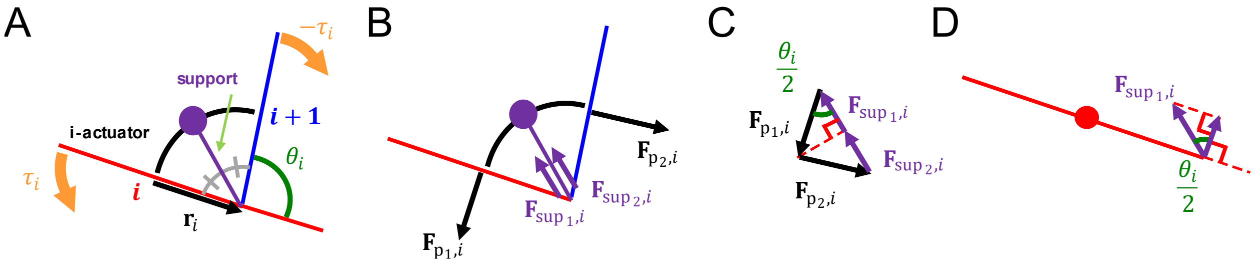


**Figure S2.** An i-actuator. (A) Composition of i-actuator. (B) Forces that i-actuator gives to i-rod and (i+1)-rod. (C) Resultant force given by i-actuator. (D) Force component of i-actuator that applies torque to i-rod.

The muscle force, which makes the torque $\tau_i$ received from i-actuator to i-rod, was designed as follows. The damped torsion spring is connected at the center of each rod and gives a force in a direction perpendicular to the rod. The support is connected to i-actuator's midpoint and i-joint (Fig. S2A). Let us say that the mass of the support and i-actuator are both 0. The support gives forces ($\mathbf{F}_{\mathrm{sup}_1,i}$, $\mathbf{F}_{\mathrm{sup}_2,i}$) of the same size to i-rod and (i+1)-rod in a direction parallel to the support (Fig. S2B). Let us assume that the resultant force received by i-actuator is $\mathbf{0}$ (Fig. S2C). Thus, the forces from the actuator generate no torque at the connection point in the middle of the rod but at the connection point at the end (Fig. S2D).

$$
\begin{aligned}
F_{\mathrm{sup}_1,i} &= F_{\mathrm{p}_1,i}\cos\frac{\theta_i}{2} \\
\tau_i &= \left(\mathbf{r}_i \times \mathbf{F}_{\mathrm{sup}_1,i}\right)\cdot\hat{\mathbf{z}} \\
&= rF_{\mathrm{sup}_1,i}\cos\frac{\theta_i}{2} \\
F_{\mathrm{sup}_1,i} &= \frac{\tau_i}{r\cos\frac{\theta_i}{2}} \\
F_{\mathrm{p}_1,i} &= \frac{F_{\mathrm{sup}_1,i}}{\cos\frac{\theta_i}{2}} \\
&= \frac{\tau_i}{r\cos^2\frac{\theta_i}{2}}
\end{aligned}
$$

The force that i-rod receives from i-actuator is $\mathbf{F}_{\mathrm{p}_1,i} + \mathbf{F}_{\mathrm{sup}_1,i}$ and the magnitude of the force is $\left|\mathbf{F}_{\mathrm{p}_1,i} + \mathbf{F}_{\mathrm{sup}_1,i}\right| = F_{\mathrm{p}_1,i}\sin(\theta_i/2) = \tau_i\sin(\theta_i/2)/\left(r\cos^2(\theta_i/2)\right)$ and the direction of the force is $-[\cos((s_i+s_{i+1})/2) \quad \sin((s_i+s_{i+1})/2)]^\mathsf{T}$. These equations come down to the following equations.

$$\mathbf{F}_{\mathrm{p}_1,i} + \mathbf{F}_{\mathrm{sup}_1,i} = -\frac{\tau_i \sin\frac{\theta_i}{2}}{r\cos^2\frac{\theta_i}{2}} \begin{bmatrix} \cos\left(\frac{s_i + s_{i+1}}{2}\right) \\ \sin\left(\frac{s_i + s_{i+1}}{2}\right) \end{bmatrix}$$

$$\mathbf{F}_{\mathrm{p}_1,i} + \mathbf{F}_{\mathrm{sup}_1,i} + \mathbf{F}_{\mathrm{p}_2,i} + \mathbf{F}_{\mathrm{sup}_2,i} = \mathbf{0}$$

$$\mathbf{F}_{\mathrm{p}_2,i} + \mathbf{F}_{\mathrm{sup}_2,i} = \frac{\tau_i \sin\frac{\theta_i}{2}}{r\cos^2\frac{\theta_i}{2}} \begin{bmatrix} \cos\left(\frac{s_i + s_{i+1}}{2}\right) \\ \sin\left(\frac{s_i + s_{i+1}}{2}\right) \end{bmatrix}$$

Therefore, the total force that i-rod receives from i-actuator and (i-1)-actuator is

$$\mathbf{F}_{c\kappa,i} = \sum_{j=-1}^{0} \frac{(-1)^{j-1} \tau_{i+j} \sin\frac{\theta_{i+j}}{2}}{r\cos^2\frac{\theta_{i+j}}{2}} \begin{bmatrix} \cos\left(\frac{s_{i+j} + s_{i+j+1}}{2}\right) \\ \sin\left(\frac{s_{i+j} + s_{i+j+1}}{2}\right) \end{bmatrix}$$

#### Joint force calculation method

Let us calculate the joint force $\mathbf{F}_i$ from the given values $(s_i, \mathbf{F}_{c\kappa,i}, \mathbf{F}_{b,i}, \tau_{c\kappa,i}, \tau_{b,i})$. In this part, the superscript $*^{\mathrm{T}}$ means the transpose of a vector or a matrix. i-rod and (i+1)-rod always meet at i-joint can be expressed as a following vector equation.

$$\mathbf{d}_i + \mathbf{r}_i = \mathbf{d}_{i+1} - \mathbf{r}_{i+1}$$

Differentiating this equation twice for time, we can see that the accelerations of the ends of i-rod and (i+1)-rod at i-joint are the same.

$$\begin{aligned}\frac{\mathrm{d}^2}{\mathrm{d}t^2}(\mathbf{d}_i + \mathbf{r}_i) &= \frac{\mathrm{d}^2}{\mathrm{d}t^2}(\mathbf{d}_i + r\hat{\mathbf{r}}_i) \\ &= \frac{\mathrm{d}}{\mathrm{d}t}\left(\mathbf{v}_i + r\omega_i\widehat{\mathbf{N}}_i\right) \\ &= \mathbf{a}_i + r\frac{\mathrm{d}\omega_i}{\mathrm{d}t}\widehat{\mathbf{N}}_i + r\omega_i\frac{d\widehat{\mathbf{N}}_i}{dt} \\ &= \mathbf{a}_i + r\alpha_i\widehat{\mathbf{N}}_i - r\omega_i^2\hat{\mathbf{r}}_i \\ &= \mathbf{a}_i + \boldsymbol{\alpha}_i \times \mathbf{r}_i - \omega_i^2\mathbf{r}_i\end{aligned}$$

$$\begin{aligned}\frac{\mathrm{d}^2}{\mathrm{d}t^2}(\mathbf{d}_{i+1} - \mathbf{r}_{i+1}) &= \frac{\mathrm{d}^2}{\mathrm{d}t^2}(\mathbf{d}_{i+1} - r\hat{\mathbf{r}}_{i+1}) \\ &= \frac{\mathrm{d}}{\mathrm{d}t}\left(\mathbf{v}_{i+1} - r\omega_{i+1}\widehat{\mathbf{N}}_{i+1}\right) \\ &= \mathbf{a}_{i+1} - r\frac{\mathrm{d}\omega_{i+1}}{\mathrm{d}t}\widehat{\mathbf{N}}_{i+1} - r\omega_{i+1}\frac{\mathrm{d}\widehat{\mathbf{N}}_{i+1}}{\mathrm{d}t} \\ &= \mathbf{a}_{i+1} - r\alpha_{i+1}\widehat{\mathbf{N}}_{i+1} + r\omega_{i+1}^2\hat{\mathbf{r}}_{i+1} \\ &= \mathbf{a}_{i+1} - \boldsymbol{\alpha}_{i+1} \times \mathbf{r}_{i+1} + \omega_{i+1}^2\mathbf{r}_{i+1}\end{aligned}$$

$$\begin{aligned}\frac{\mathrm{d}^2}{\mathrm{d}t^2}(\mathbf{d}_i + \mathbf{r}_i) &= \frac{\mathrm{d}^2}{\mathrm{d}t^2}(\mathbf{d}_{i+1} - \mathbf{r}_{i+1}) \\ &= \mathbf{a}_i + \boldsymbol{\alpha}_i \times \mathbf{r}_i - \omega_i^2\mathbf{r}_i \\ &= \mathbf{a}_{i+1} - \boldsymbol{\alpha}_{i+1} \times \mathbf{r}_{i+1} + \omega_{i+1}^2\mathbf{r}_{i+1}\end{aligned}$$

Multiplying both sides by $m$ gives:

$$m(\mathbf{a}_i + \boldsymbol{\alpha}_i \times \mathbf{r}_i) - m\omega_i^2\mathbf{r}_i = m(\mathbf{a}_{i+1} - \boldsymbol{\alpha}_{i+1} \times \mathbf{r}_{i+1}) + m\omega_{i+1}^2\mathbf{r}_{i+1}$$

If $\mathbf{F}_{\mathrm{res},i}$ is the total force applied to i-rod other than $\mathbf{F}_i$ and $-\mathbf{F}_{i-1}$ , which is $\mathbf{F}_{\mathrm{res},i} = \mathbf{F}_{c\kappa,i} + \mathbf{F}_{b,i}$, then:

$$m\mathbf{a}_i = \mathbf{F}_i - \mathbf{F}_{i-1} + \mathbf{F}_{\mathrm{res},i}$$

If $\boldsymbol{\tau}_{\mathrm{res},i}$ is the total torque applied to i-rod excluding $\mathbf{r}_i \times (\mathbf{F}_{i-1} + \mathbf{F}_i)$ , which is $\boldsymbol{\tau}_{\mathrm{res},i} = \tau_{c\kappa,i} + \tau_{b,i}$ , then:

$$I\boldsymbol{\alpha}_i = \frac{1}{3}mr^2\boldsymbol{\alpha}_i = \mathbf{r}_i \times (\mathbf{F}_{i-1} + \mathbf{F}_i) + \boldsymbol{\tau}_{\text{res},i}$$

Dividing both sides by $\frac{1}{3}r^2$ gives:

$$m\boldsymbol{\alpha}_i = 3r^{-2}\mathbf{r}_i \times (\mathbf{F}_{i-1} + \mathbf{F}_i) + 3r^{-2}\boldsymbol{\tau}_{\text{res},i}$$

If $\mathbf{h}_{\text{res},i} \equiv 3r^{-2}\boldsymbol{\tau}_{\text{res},i} \times \mathbf{r}_i$ , then:

$$\begin{aligned} m(\mathbf{a}_i + \boldsymbol{\alpha}_i \times \mathbf{r}_i) &= \left(\mathbf{F}_i - \mathbf{F}_{i-1} + \mathbf{F}_{\text{res},i}\right) + \left[3r^{-2}\mathbf{r}_i \times (\mathbf{F}_{i-1} + \mathbf{F}_i) + 3r^{-2}\boldsymbol{\tau}_{\text{res},i}\right] \times \mathbf{r}_i \\ &= (\mathbf{F}_i - \mathbf{F}_{i-1}) + 3[\hat{\mathbf{r}}_i \times (\mathbf{F}_{i-1} + \mathbf{F}_i)] \times \hat{\mathbf{r}}_i + \mathbf{F}_{\text{res},i} + 3r^{-2}\boldsymbol{\tau}_{\text{res},i} \times \mathbf{r}_i \\ &= (\mathbf{F}_i - \mathbf{F}_{i-1}) + 3\left[(\mathbf{F}_{i-1} + \mathbf{F}_i) \cdot \widehat{\mathbf{N}}_i\right]\widehat{\mathbf{N}}_i + \mathbf{F}_{\text{res},i} + \mathbf{h}_{\text{res},i} \end{aligned}$$

Following the same method,

$$m(\mathbf{a}_{i+1} - \boldsymbol{\alpha}_{i+1} \times \mathbf{r}_{i+1}) = (\mathbf{F}_{i+1} - \mathbf{F}_i) - 3\left[(\mathbf{F}_i + \mathbf{F}_{i+1}) \cdot \widehat{\mathbf{N}}_{i+1}\right]\widehat{\mathbf{N}}_{i+1} + \mathbf{F}_{\text{res},i+1} - \mathbf{h}_{\text{res},i+1}$$

To organize the terms of the equations into known values and unknown values,

$$\begin{aligned} \mathbf{k}_i &\equiv \mathbf{F}_i - \mathbf{F}_{i-1} \\ \mathbf{P}_i &\equiv 3\widehat{\mathbf{N}}_i\widehat{\mathbf{N}}_i^{\top} \\ \mathbf{h}_i &\equiv 3\left[(\mathbf{F}_{i-1} + \mathbf{F}_i) \cdot \widehat{\mathbf{N}}_i\right]\widehat{\mathbf{N}}_i \\ &= 3\widehat{\mathbf{N}}_i\widehat{\mathbf{N}}_i^{\top}(\mathbf{F}_{i-1} + \mathbf{F}_i) \\ &= \mathbf{P}_i(\mathbf{F}_{i-1} + \mathbf{F}_i) \\ m(\mathbf{a}_i + \boldsymbol{\alpha}_i \times \mathbf{r}_i) &= \mathbf{k}_i + \mathbf{h}_i + \mathbf{F}_{\text{res},i} + \mathbf{h}_{\text{res},i} \\ m(\mathbf{a}_{i+1} - \boldsymbol{\alpha}_{i+1} \times \mathbf{r}_{i+1}) &= \mathbf{k}_{i+1} - \mathbf{h}_{i+1} + \mathbf{F}_{\text{res},i+1} - \mathbf{h}_{\text{res},i+1} \\ \mathbf{k}_i + \mathbf{h}_i + \mathbf{F}_{\text{res},i} + \mathbf{h}_{\text{res},i} - m\omega_i^2\mathbf{r}_i &= \mathbf{k}_{i+1} - \mathbf{h}_{i+1} + \mathbf{F}_{\text{res},i+1} - \mathbf{h}_{\text{res},i+1} + m\omega_{i+1}^2\mathbf{r}_{i+1} \\ \mathbf{q}_i &\equiv \mathbf{k}_i + \mathbf{h}_i - \mathbf{k}_{i+1} + \mathbf{h}_{i+1} \\ &= -\mathbf{F}_{\text{res},i} - \mathbf{h}_{\text{res},i} + \mathbf{F}_{\text{res},i+1} - \mathbf{h}_{\text{res},i+1} + m\omega_i^2\mathbf{r}_i + m\omega_{i+1}^2\mathbf{r}_{i+1} \end{aligned}$$

Therefore, if we know all $\mathbf{F}_{\text{res},i}$ and $\boldsymbol{\tau}_{\text{res},i}$ for each i, we can find $\mathbf{q}_i$.

If $\mathbf{I} \equiv \begin{bmatrix} 1 & 0 \\ 0 & 1 \end{bmatrix}$, and if we expand $\mathbf{q}_i$,

$$\begin{aligned} \mathbf{q}_i \equiv\ & \mathbf{k}_i + \mathbf{h}_i - \mathbf{k}_{i+1} + \mathbf{h}_{i+1} \\ =\ & -\mathbf{k}_{i+1} + \mathbf{k}_i + \mathbf{h}_i + \mathbf{h}_{i+1} \\ =\ & -\mathbf{F}_{i+1} + \mathbf{F}_i + \mathbf{F}_i - \mathbf{F}_{i-1} + \mathbf{P}_i(\mathbf{F}_{i-1} + \mathbf{F}_i) + \mathbf{P}_{i+1}(\mathbf{F}_i + \mathbf{F}_{i+1}) \\ =\ & \mathbf{F}_{i-1}(\mathbf{P}_i - \mathbf{I}) + \mathbf{F}_i(\mathbf{P}_i + \mathbf{P}_{i+1} + 2\mathbf{I}) + \mathbf{F}_{i+1}(\mathbf{P}_{i+1} - \mathbf{I}) \end{aligned}$$

If we set $\mathbf{A}_i \equiv \mathbf{P}_i - \mathbf{I}$ and $\mathbf{B}_i \equiv \mathbf{P}_i + \mathbf{P}_{i+1} + 2\mathbf{I}$, then:

$$\mathbf{q}_i = \mathbf{A}_i\mathbf{F}_{i-1} + \mathbf{B}_i\mathbf{F}_i + \mathbf{A}_{i+1}\mathbf{F}_{i+1}$$

If we set $\overline{\mathbf{0}} = \begin{bmatrix} 0 & 0 \\ 0 & 0 \end{bmatrix}$ and express the above equation in a multidimensional tensor form,

$$\begin{bmatrix} \mathbf{B}_1 & \mathbf{A}_2 & \overline{\mathbf{0}} & \overline{\mathbf{0}} & \overline{\mathbf{0}} & \cdots & \overline{\mathbf{0}} \\ \mathbf{A}_2 & \mathbf{B}_2 & \mathbf{A}_3 & \overline{\mathbf{0}} & \overline{\mathbf{0}} & \cdots & \overline{\mathbf{0}} \\ \overline{\mathbf{0}} & \mathbf{A}_3 & \mathbf{B}_3 & \mathbf{A}_4 & \overline{\mathbf{0}} & \cdots & \overline{\mathbf{0}} \\ \overline{\mathbf{0}} & \overline{\mathbf{0}} & \mathbf{A}_4 & \mathbf{B}_4 & \mathbf{A}_5 & \cdots & \overline{\mathbf{0}} \\ \overline{\mathbf{0}} & \overline{\mathbf{0}} & \overline{\mathbf{0}} & \mathbf{A}_5 & \mathbf{B}_5 & \ddots & \vdots \\ \vdots & \vdots & \vdots & \vdots & \ddots & \ddots & \mathbf{A}_{n-1} \\ \overline{\mathbf{0}} & \overline{\mathbf{0}} & \overline{\mathbf{0}} & \overline{\mathbf{0}} & \cdots & \mathbf{A}_{n-1} & \mathbf{B}_{n-1} \end{bmatrix} \begin{bmatrix} \mathbf{F}_1 \\ \mathbf{F}_2 \\ \mathbf{F}_3 \\ \mathbf{F}_4 \\ \vdots \\ \mathbf{F}_{n-2} \\ \mathbf{F}_{n-1} \end{bmatrix} = \begin{bmatrix} \mathbf{q}_1 \\ \mathbf{q}_2 \\ \mathbf{q}_3 \\ \mathbf{q}_4 \\ \vdots \\ \mathbf{q}_{n-2} \\ \mathbf{q}_{n-1} \end{bmatrix}$$

Let us set:

$$\mathcal{D} \equiv \begin{bmatrix} \mathbf{B}_1 & \mathbf{A}_2 & \overline{\mathbf{0}} & \overline{\mathbf{0}} & \overline{\mathbf{0}} & \cdots & \overline{\mathbf{0}} \\ \mathbf{A}_2 & \mathbf{B}_2 & \mathbf{A}_3 & \overline{\mathbf{0}} & \overline{\mathbf{0}} & \cdots & \overline{\mathbf{0}} \\ \overline{\mathbf{0}} & \mathbf{A}_3 & \mathbf{B}_3 & \mathbf{A}_4 & \overline{\mathbf{0}} & \cdots & \overline{\mathbf{0}} \\ \overline{\mathbf{0}} & \overline{\mathbf{0}} & \mathbf{A}_4 & \mathbf{B}_4 & \mathbf{A}_5 & \cdots & \overline{\mathbf{0}} \\ \overline{\mathbf{0}} & \overline{\mathbf{0}} & \overline{\mathbf{0}} & \mathbf{A}_5 & \mathbf{B}_5 & \ddots & \vdots \\ \vdots & \vdots & \vdots & \vdots & \ddots & \ddots & \mathbf{A}_{n-1} \\ \overline{\mathbf{0}} & \overline{\mathbf{0}} & \overline{\mathbf{0}} & \overline{\mathbf{0}} & \cdots & \mathbf{A}_{n-1} & \mathbf{B}_{n-1} \end{bmatrix}$$

$$\mathcal{F} \equiv \begin{bmatrix} \mathbf{F}_1 \\ \mathbf{F}_2 \\ \mathbf{F}_3 \\ \mathbf{F}_4 \\ \vdots \\ \mathbf{F}_{n-2} \\ \mathbf{F}_{n-1} \end{bmatrix}$$

$$\mathcal{Q} \equiv \begin{bmatrix} \mathbf{q}_1 \\ \mathbf{q}_2 \\ \mathbf{q}_3 \\ \mathbf{q}_4 \\ \vdots \\ \mathbf{q}_{n-2} \\ \mathbf{q}_{n-1} \end{bmatrix}$$

As $\mathbf{A}_i$ and $\mathbf{B}_i$ can be calculated from $\mathbf{P}_i$, $\mathbf{P}_i$ from $\widehat{\mathbf{N}}_i$, and $\widehat{\mathbf{N}}_i$ from $s_i$, we can find $\mathcal{D}$ from $s_i$.

$$\begin{aligned} \mathbf{q}_i = & \ -\mathbf{F}_{\text{res},i} + \mathbf{F}_{\text{res},i+1} - \mathbf{h}_{\text{res},i} - \mathbf{h}_{\text{res},i+1} + m\omega_i^2\mathbf{r}_i + m\omega_{i+1}^2\mathbf{r}_{i+1} \\ = & \ -\mathbf{F}_{\text{res},i} + \mathbf{F}_{\text{res},i+1} - 3r^{-2}\left(\boldsymbol{\tau}_{\text{res},i} \times \mathbf{r}_i + \boldsymbol{\tau}_{\text{res},i+1} \times \mathbf{r}_{i+1}\right) + m\omega_i^2\mathbf{r}_i + m\omega_{i+1}^2\mathbf{r}_{i+1} \end{aligned}$$

Therefore, if we know $\mathbf{F}_{c\kappa,i}$, $\mathbf{F}_{b,i}$, $\tau_{c\kappa,i}$, $\tau_{b,i}$ we can find $\mathcal{Q}$.

$$\therefore \mathcal{D}\mathcal{F} = \mathcal{Q}$$

We can find $\mathcal{F}$, thus $\mathbf{F}_i$ $(i \in \{1, \cdots, n-1\})$, by solving this tensor equation.

If we denote each component of the tensor, which are each matrix and the p-th row q-th column component of the vector, as $(*)_{p,q}$, then:

$$\begin{bmatrix} \begin{bmatrix} (\mathbf{B}_1)_{1,1} & (\mathbf{B}_1)_{1,2} \\ (\mathbf{B}_1)_{2,1} & (\mathbf{B}_1)_{2,2} \end{bmatrix} & \begin{bmatrix} (\mathbf{A}_2)_{1,1} & (\mathbf{A}_2)_{1,2} \\ (\mathbf{A}_2)_{2,1} & (\mathbf{A}_2)_{2,2} \end{bmatrix} & \begin{bmatrix} 0 & 0 \\ 0 & 0 \end{bmatrix} & \cdots \\ \begin{bmatrix} (\mathbf{A}_2)_{1,1} & (\mathbf{A}_2)_{1,2} \\ (\mathbf{A}_2)_{2,1} & (\mathbf{A}_2)_{2,2} \end{bmatrix} & \begin{bmatrix} (\mathbf{B}_2)_{1,1} & (\mathbf{B}_2)_{1,2} \\ (\mathbf{B}_2)_{2,1} & (\mathbf{B}_2)_{2,2} \end{bmatrix} & \begin{bmatrix} (\mathbf{A}_3)_{1,1} & (\mathbf{A}_3)_{1,2} \\ (\mathbf{A}_3)_{2,1} & (\mathbf{A}_3)_{2,2} \end{bmatrix} & \cdots \\ \begin{bmatrix} 0 & 0 \\ 0 & 0 \end{bmatrix} & \begin{bmatrix} (\mathbf{A}_3)_{1,1} & (\mathbf{A}_3)_{1,2} \\ (\mathbf{A}_3)_{2,1} & (\mathbf{A}_3)_{2,2} \end{bmatrix} & \begin{bmatrix} (\mathbf{B}_3)_{1,1} & (\mathbf{B}_3)_{1,2} \\ (\mathbf{B}_3)_{2,1} & (\mathbf{B}_3)_{2,2} \end{bmatrix} & \cdots \\ \vdots & \vdots & \vdots & \ddots \end{bmatrix} \begin{bmatrix} \begin{bmatrix} (\mathbf{F}_1)_{1,1} \\ (\mathbf{F}_1)_{2,1} \end{bmatrix} \\ \begin{bmatrix} (\mathbf{F}_2)_{1,1} \\ (\mathbf{F}_2)_{2,1} \end{bmatrix} \\ \begin{bmatrix} (\mathbf{F}_3)_{1,1} \\ (\mathbf{F}_3)_{2,1} \end{bmatrix} \\ \vdots \end{bmatrix} = \begin{bmatrix} \begin{bmatrix} (\mathbf{q}_1)_{1,1} \\ (\mathbf{q}_1)_{2,1} \end{bmatrix} \\ \begin{bmatrix} (\mathbf{q}_2)_{1,1} \\ (\mathbf{q}_2)_{2,1} \end{bmatrix} \\ \begin{bmatrix} (\mathbf{q}_3)_{1,1} \\ (\mathbf{q}_3)_{2,1} \end{bmatrix} \\ \vdots \end{bmatrix}$$

And the matrix equation equivalent to this tensor equation is:

$$\begin{bmatrix} (\mathbf{B}_1)_{1,1} & (\mathbf{B}_1)_{1,2} & (\mathbf{A}_2)_{1,1} & (\mathbf{A}_2)_{1,2} & 0 & 0 & \cdots \\ (\mathbf{B}_1)_{2,1} & (\mathbf{B}_1)_{2,2} & (\mathbf{A}_2)_{2,1} & (\mathbf{A}_2)_{2,2} & 0 & 0 & \cdots \\ (\mathbf{A}_2)_{1,1} & (\mathbf{A}_2)_{1,2} & (\mathbf{B}_2)_{1,1} & (\mathbf{B}_2)_{1,2} & (\mathbf{A}_3)_{1,1} & (\mathbf{A}_3)_{1,2} & \cdots \\ (\mathbf{A}_2)_{2,1} & (\mathbf{A}_2)_{2,2} & (\mathbf{B}_2)_{2,1} & (\mathbf{B}_2)_{2,2} & (\mathbf{A}_3)_{2,1} & (\mathbf{A}_3)_{2,2} & \cdots \\ 0 & 0 & (\mathbf{A}_3)_{1,1} & (\mathbf{A}_3)_{1,2} & (\mathbf{B}_3)_{1,1} & (\mathbf{B}_3)_{1,2} & \cdots \\ 0 & 0 & (\mathbf{A}_3)_{2,1} & (\mathbf{A}_3)_{2,2} & (\mathbf{B}_3)_{2,1} & (\mathbf{B}_3)_{2,2} & \cdots \\ \vdots & \vdots & \vdots & \vdots & \vdots & \vdots & \ddots \end{bmatrix} \begin{bmatrix} (\mathbf{F}_1)_{1,1} \\ (\mathbf{F}_1)_{2,1} \\ (\mathbf{F}_2)_{1,1} \\ (\mathbf{F}_2)_{2,1} \\ (\mathbf{F}_3)_{1,1} \\ (\mathbf{F}_3)_{2,1} \\ \vdots \end{bmatrix} = \begin{bmatrix} (\mathbf{q}_1)_{1,1} \\ (\mathbf{q}_1)_{2,1} \\ (\mathbf{q}_2)_{1,1} \\ (\mathbf{q}_2)_{2,1} \\ (\mathbf{q}_3)_{1,1} \\ (\mathbf{q}_3)_{2,1} \\ \vdots \end{bmatrix}$$

Let us set:

$$\mathbf{D} \equiv \begin{bmatrix} (\mathbf{B}_1)_{1,1} & (\mathbf{B}_1)_{1,2} & (\mathbf{A}_2)_{1,1} & (\mathbf{A}_2)_{1,2} & 0 & 0 & \cdots \\ (\mathbf{B}_1)_{2,1} & (\mathbf{B}_1)_{2,2} & (\mathbf{A}_2)_{2,1} & (\mathbf{A}_2)_{2,2} & 0 & 0 & \cdots \\ (\mathbf{A}_2)_{1,1} & (\mathbf{A}_2)_{1,2} & (\mathbf{B}_2)_{1,1} & (\mathbf{B}_2)_{1,2} & (\mathbf{A}_3)_{1,1} & (\mathbf{A}_3)_{1,2} & \cdots \\ (\mathbf{A}_2)_{2,1} & (\mathbf{A}_2)_{2,2} & (\mathbf{B}_2)_{2,1} & (\mathbf{B}_2)_{2,2} & (\mathbf{A}_3)_{2,1} & (\mathbf{A}_3)_{2,2} & \cdots \\ 0 & 0 & (\mathbf{A}_3)_{1,1} & (\mathbf{A}_3)_{1,2} & (\mathbf{B}_3)_{1,1} & (\mathbf{B}_3)_{1,2} & \cdots \\ 0 & 0 & (\mathbf{A}_3)_{2,1} & (\mathbf{A}_3)_{2,2} & (\mathbf{B}_3)_{2,1} & (\mathbf{B}_3)_{2,2} & \cdots \\ \vdots & \vdots & \vdots & \vdots & \vdots & \vdots & \ddots \end{bmatrix}$$

Since $\mathbf{A}_i$, $\mathbf{B}_i$ are symmetric matrices, $\mathbf{D}$ is a heptadiagonal symmetric matrix. Since $\mathbf{D}$ is a symmetric matrix, the solution to the matrix equation can be found with the Cholesky decomposition. Therefore, we can find the x-axis and y-axis components of $\mathbf{F}_i$.

As a result, we can find the joint force $\mathbf{F}_i$ from the known values ($s_i$, $\mathbf{F}_{c\kappa,i}$, $\mathbf{F}_{b,i}$, $\tau_{c\kappa,i}$, $\tau_{b,i}$).

#### Proof of numerical integration for the translational motion of a worm using semi-implicit Euler method

When the friction coefficients $b_{\perp}$ , $b_{\parallel}$ are sufficiently large compared to $M/\Delta t$, numerical integration via the explicit Euler method $\mathbf{v}_{\mathrm{c}}^{(t+\Delta t)} = \mathbf{v}_{\mathrm{c}}^{(t)} + \mathbf{a}_{\mathrm{c}}^{(t)}\Delta t = \mathbf{v}_{\mathrm{c}}^{(t)} + \frac{\sum_i \mathbf{F}_{b,i}^{(t)}}{M}\Delta t)$ becomes unstable(Butcher, 2004). Therefore, for all friction coefficients greater than or equal to 0, the semi-implicit Euler method $\mathbf{v}_{\mathrm{c}}^{(t+\Delta t)} = \mathbf{v}_{\mathrm{c}}^{(t)} + \mathbf{a}_{\mathrm{c}}^{(t+\Delta t)}\Delta t \simeq \mathbf{v}_{\mathrm{c}}^{(t)} + \frac{1}{1+\frac{b_{\perp}\Delta t}{M}}\frac{\sum_i \mathbf{F}_{b,i}^{(t)}}{M}\Delta t)$ was used to ensure numerical integration remains stable, and its proof is as follows.

Newton's equation for the translational motion of each i-rod is as follows.

$$\begin{aligned}
\mathbf{F}_i &= m\mathbf{a}_i \\
&= \mathbf{F}_{b,i} + \mathbf{F}_{c\kappa,i} + \mathbf{F}_{\mathrm{joint},i} \\
&= -\frac{b_{\perp}}{n}\mathbf{v}_{\perp,i} - \frac{b_{\parallel}}{n}\mathbf{v}_{\parallel,i} + \mathbf{F}_{c\kappa,i} + \mathbf{F}_{\mathrm{joint},i}
\end{aligned}$$

The integration formula for $\mathbf{a}_i$ using the Implicit Euler Method is as follows. (where $b_{\perp} \geq b_{\parallel}$)

$$\begin{aligned}
\mathbf{v}_i^{(t+\Delta t)} &= \mathbf{v}_i^{(t)} + \mathbf{a}_i^{(t+\Delta t)}\Delta t \\
&= \mathbf{v}_i^{(t)} - \frac{\Delta t}{m}\left(\frac{b_{\perp}}{n}\mathbf{v}_{\perp,i}^{(t+\Delta t)} + \frac{b_{\parallel}}{n}\mathbf{v}_{\parallel,i}^{(t+\Delta t)}\right) + \frac{\Delta t}{m}\left(\mathbf{F}_{c\kappa,i}^{(t+\Delta t)} + \mathbf{F}_{\mathrm{joint},i}^{(t+\Delta t)}\right) \\
&= \mathbf{v}_i^{(t)} - \frac{\Delta t}{M}\left(b_{\perp}\mathbf{v}_{\perp,i}^{(t+\Delta t)} + b_{\parallel}\mathbf{v}_{\parallel,i}^{(t+\Delta t)}\right) + \frac{\Delta t}{m}\left(\mathbf{F}_{c\kappa,i}^{(t+\Delta t)} + \mathbf{F}_{\mathrm{joint},i}^{(t+\Delta t)}\right) \\
&= \mathbf{v}_i^{(t)} - \frac{\Delta t}{M}\left(b_{\perp}\left(\mathbf{v}_{\perp,i}^{(t+\Delta t)} + \mathbf{v}_{\parallel,i}^{(t+\Delta t)}\right) + (b_{\parallel} - b_{\perp})\mathbf{v}_{\parallel,i}^{(t+\Delta t)}\right) + \frac{\Delta t}{m}\left(\mathbf{F}_{c\kappa,i}^{(t+\Delta t)} + \mathbf{F}_{\mathrm{joint},i}^{(t+\Delta t)}\right) \\
&= \mathbf{v}_i^{(t)} - \frac{\Delta t}{M}\left(b_{\perp}\mathbf{v}_i^{(t+\Delta t)} + (b_{\parallel} - b_{\perp})\mathbf{v}_{\parallel,i}^{(t+\Delta t)}\right) + \frac{\Delta t}{m}\left(\mathbf{F}_{c\kappa,i}^{(t+\Delta t)} + \mathbf{F}_{\mathrm{joint},i}^{(t+\Delta t)}\right) \\
\left(1 + \frac{b_{\perp}\Delta t}{M}\right)\mathbf{v}_i^{(t+\Delta t)} &= \mathbf{v}_i^{(t)} - \frac{\Delta t}{M}\left((b_{\parallel} - b_{\perp})\mathbf{v}_{\parallel,i}^{(t+\Delta t)}\right) + \frac{\Delta t}{m}\left(\mathbf{F}_{c\kappa,i}^{(t+\Delta t)} + \mathbf{F}_{\mathrm{joint},i}^{(t+\Delta t)}\right) \\
&= \mathbf{v}_i^{(t)} + \frac{(b_{\perp} - b_{\parallel})\Delta t}{M}\mathbf{v}_{\parallel,i}^{(t+\Delta t)} + \frac{\Delta t}{m}\left(\mathbf{F}_{c\kappa,i}^{(t+\Delta t)} + \mathbf{F}_{\mathrm{joint},i}^{(t+\Delta t)}\right)
\end{aligned}$$

Because $\mathbf{v}_{\perp,i}^{(t+\Delta t)}$ and $\mathbf{v}_{\parallel,i}^{(t+\Delta t)}$ are unknown at time t, the numerical calculation of the above formula is impossible.

$$\frac{\left|\frac{(b_{\perp} - b_{\parallel})\Delta t}{M}\left(\mathbf{v}_{\parallel,i}^{(t+\Delta t)} - \mathbf{v}_{\parallel,i}^{(t)}\right)\right|}{\left|\mathbf{v}_i^{(t)} + \frac{(b_{\perp} - b_{\parallel})\Delta t}{M}\mathbf{v}_{\parallel,i}^{(t+\Delta t)} + \frac{\Delta t}{m}\left(\mathbf{F}_{c\kappa,i}^{(t+\Delta t)} + \mathbf{F}_{\mathrm{joint},i}^{(t+\Delta t)}\right)\right|} \simeq 0$$

If the above formula is assumed to be true, the following approximation can be used.

$$\begin{aligned}\left(1+\frac{b_\perp \Delta t}{M}\right)\mathbf{v}_i^{(t+\Delta t)} &= \mathbf{v}_i^{(t)} + \frac{(b_\perp - b_\parallel)\Delta t}{M}\mathbf{v}_{\parallel,i}^{(t+\Delta t)} + \frac{\Delta t}{m}\left(\mathbf{F}_{c\kappa,i}^{(t+\Delta t)} + \mathbf{F}_{\mathrm{joint},i}^{(t+\Delta t)}\right)\\ &\simeq \mathbf{v}_i^{(t)} + \frac{(b_\perp - b_\parallel)\Delta t}{M}\mathbf{v}_{\parallel,i}^{(t)} + \frac{\Delta t}{m}\left(\mathbf{F}_{c\kappa,i}^{(t+\Delta t)} + \mathbf{F}_{\mathrm{joint},i}^{(t+\Delta t)}\right)\end{aligned}$$

The approximate value of $\mathbf{v}_i^{(t+\Delta t)}$ by the above approximation is as follows.

$$\mathbf{v}_i^{(t+\Delta t)} \simeq \left[\mathbf{v}_i^{(t)} + \frac{(b_\perp - b_\parallel)\Delta t}{M}\mathbf{v}_{\parallel,i}^{(t)} + \frac{\Delta t}{m}\left(\mathbf{F}_{c\kappa,i}^{(t+\Delta t)} + \mathbf{F}_{\mathrm{joint},i}^{(t+\Delta t)}\right)\right] \Big/ \left(1+\frac{b_\perp \Delta t}{M}\right)$$

The approximate value of $\mathbf{a}_i^{(t+\Delta t)}\Delta t$ by the approximate value of $\mathbf{v}_i^{(t+\Delta t)}$ is as follows.

$$\begin{aligned}\mathbf{a}_i^{(t+\Delta t)}\Delta t &= \mathbf{v}_i^{(t+\Delta t)} - \mathbf{v}_i^{(t)}\\ &\simeq \left[\mathbf{v}_i^{(t)} + \frac{(b_\perp - b_\parallel)\Delta t}{M}\mathbf{v}_{\parallel,i}^{(t)} + \frac{\Delta t}{m}\left(\mathbf{F}_{c\kappa,i}^{(t+\Delta t)} + \mathbf{F}_{\mathrm{joint},i}^{(t+\Delta t)}\right)\right] \Big/ \left(1+\frac{b_\perp \Delta t}{M}\right) - \mathbf{v}_i^{(t)}\\ &= \left[-\frac{b_\perp \Delta t}{M}\mathbf{v}_i^{(t)} + \frac{(b_\perp - b_\parallel)\Delta t}{M}\mathbf{v}_{\parallel,i}^{(t)} + \frac{\Delta t}{m}\left(\mathbf{F}_{c\kappa,i}^{(t+\Delta t)} + \mathbf{F}_{\mathrm{joint},i}^{(t+\Delta t)}\right)\right] \Big/ \left(1+\frac{b_\perp \Delta t}{M}\right)\\ &= \frac{\Delta t}{m}\left[-\frac{b_\perp}{n}\mathbf{v}_i^{(t)} + \frac{(b_\perp - b_\parallel)}{n}\mathbf{v}_{\parallel,i}^{(t)} + \mathbf{F}_{c\kappa,i}^{(t+\Delta t)} + \mathbf{F}_{\mathrm{joint},i}^{(t+\Delta t)}\right] \Big/ \left(1+\frac{b_\perp \Delta t}{M}\right)\\ &= \frac{\Delta t}{m}\left[-\frac{b_\perp}{n}\left(\mathbf{v}_i^{(t)} - \mathbf{v}_{\parallel,i}^{(t)}\right) - \frac{b_\parallel}{n}\mathbf{v}_{\parallel,i}^{(t)} + \mathbf{F}_{c\kappa,i}^{(t+\Delta t)} + \mathbf{F}_{\mathrm{joint},i}^{(t+\Delta t)}\right] \Big/ \left(1+\frac{b_\perp \Delta t}{M}\right)\\ &= \frac{\Delta t}{m}\left[-\frac{b_\perp}{n}\mathbf{v}_{\perp,i}^{(t)} - \frac{b_\parallel}{n}\mathbf{v}_{\parallel,i}^{(t)} + \mathbf{F}_{c\kappa,i}^{(t+\Delta t)} + \mathbf{F}_{\mathrm{joint},i}^{(t+\Delta t)}\right] \Big/ \left(1+\frac{b_\perp \Delta t}{M}\right)\\ &= \frac{\Delta t}{m}\left[\mathbf{F}_{b,i}^{(t)} + \mathbf{F}_{c\kappa,i}^{(t+\Delta t)} + \mathbf{F}_{\mathrm{joint},i}^{(t+\Delta t)}\right] \Big/ \left(1+\frac{b_\perp \Delta t}{M}\right)\end{aligned}$$

If both sides are divided by $\Delta t$,

$$\mathbf{a}_i^{(t+\Delta t)} \simeq \frac{1}{\left(1+\frac{b_\perp \Delta t}{M}\right)}\frac{\mathbf{F}_{b,i}^{(t)} + \mathbf{F}_{c\kappa,i}^{(t+\Delta t)} + \mathbf{F}_{\mathrm{joint},i}^{(t+\Delta t)}}{m}$$

$\mathbf{F}_{c\kappa,i}^{(t+\Delta t)}$, $\mathbf{F}_{\mathrm{joint},i}^{(t+\Delta t)}$ satisfy the following because they are the internal forces of the worm ($\vec{0}$ is a zero vector).

$$\begin{aligned}\sum_i \mathbf{F}_{c\kappa,i}^{(t+\Delta t)} &= \vec{0}\\ \sum_i \mathbf{F}_{\mathrm{joint},i}^{(t+\Delta t)} &= \vec{0}\end{aligned}$$

Therefore, the approximation of the force $M\mathbf{a}_\mathrm{c}^{(t+\Delta t)}$ received by the worm is as follows.

$$
\begin{aligned}
M\mathbf{a}_{\mathrm{c}}^{(t+\Delta t)} &= \sum_i m\,\mathbf{a}_i^{(t+\Delta t)} \\
&\simeq \sum_i m \frac{1}{\left(1+\frac{b_{\perp}\Delta t}{M}\right)} \frac{\mathbf{F}_{b,i}^{(t)} + \mathbf{F}_{c\kappa,i}^{(t+\Delta t)} + \mathbf{F}_{\mathrm{joint},i}^{(t+\Delta t)}}{m} \\
&= \frac{1}{\left(1+\frac{b_{\perp}\Delta t}{M}\right)} \left( \sum_i \mathbf{F}_{b,i}^{(t)} + \sum_i \mathbf{F}_{c\kappa,i}^{(t+\Delta t)} + \sum_i \mathbf{F}_{\mathrm{joint},i}^{(t+\Delta t)} \right) \\
&= \frac{1}{\left(1+\frac{b_{\perp}\Delta t}{M}\right)} \sum_i \mathbf{F}_{b,i}^{(t)}
\end{aligned}
$$

If both sides are divided by $M$,

$$
\mathbf{a}_{\mathrm{c}}^{(t+\Delta t)} \simeq \frac{1}{\left(1+\frac{b_{\perp}\Delta t}{M}\right)} \frac{\sum_i \mathbf{F}_{b,i}^{(t)}}{M}
$$

This approximation ensures computational stability regardless of the size of $b_{\perp}$ , $b_{\parallel}$. That is, this approximation solves the problem of the decrease in computational stability of numerical integration through the explicit Euler method when $b_{\perp}$ , $b_{\parallel}$ are sufficiently large compared to $M/\Delta t$.

#### Numerical integration of the rotational motion of i-rod using semi-implicit Euler method

First, the numerical integration formulas for $\omega_i$ and $\alpha_i$ using the Implicit Euler method are as follows.

$$s_i^{(t+\Delta t)} = s_i^{(t)} + \omega_i^{(t+\Delta t)}\Delta t$$

$$\omega_i^{(t+\Delta t)} = \omega_i^{(t)} + \alpha_i^{(t+\Delta t)}\Delta t$$

If we set $\beta \equiv \frac{1}{3}\frac{b_\perp}{n}r^2$, the equation describing the rotation of i-rod is as follows.

$$\begin{aligned}
I\alpha_i &= \tau_{b,i} + \tau_{c\kappa,i} + \tau_{\text{joint},i} \\
&= -\beta\omega_i + c(\omega_{i+1} - 2\omega_i + \omega_{i-1}) + \kappa(\theta_i - \theta_{i-1} - \theta_{\text{ctrl},i} + \theta_{\text{ctrl},i-1}) + \tau_{\text{joint},i} \\
&= -\beta\omega_i + c(\omega_{i+1} - 2\omega_i + \omega_{i-1}) + \kappa(s_{i+1} - 2s_i + s_{i-1} - \theta_{\text{ctrl},i} + \theta_{\text{ctrl},i-1}) + \tau_{\text{joint},i} \\
&= c\omega_{i-1} - (\beta + 2c)\omega_i + c\omega_{i+1} + \kappa(s_{i-1} - 2s_i + s_{i+1}) - \kappa(\theta_{\text{ctrl},i} - \theta_{\text{ctrl},i+1}) + \tau_{\text{joint},i}
\end{aligned}$$

If the above formula is expanded for time $t + \Delta t$,

$$\begin{aligned}
I\alpha_i^{(t+\Delta t)} = &\ c\omega_{i-1}^{(t+\Delta t)} - (\beta + 2c)\omega_i^{(t+\Delta t)} + c\omega_{i+1}^{(t+\Delta t)} \\
&+\kappa(s_{i-1}^{(t+\Delta t)} - 2s_i^{(t+\Delta t)} + s_{i+1}^{(t+\Delta t)}) - \kappa(\theta_{\text{ctrl},i}^{(t+\Delta t)} - \theta_{\text{ctrl},i+1}^{(t+\Delta t)}) + \tau_{\text{joint},i}^{(t+\Delta t)} \\
= &\ c\omega_{i-1}^{(t+\Delta t)} - (\beta + 2c)\omega_i^{(t+\Delta t)} + c\omega_{i+1}^{(t+\Delta t)} \\
&+\kappa\left((s_{i-1}^{(t)} + \omega_{i-1}^{(t+\Delta t)}\Delta t) - 2(s_i^{(t)} + \omega_i^{(t+\Delta t)}\Delta t) + (s_{i+1}^{(t)} + \omega_{i+1}^{(t+\Delta t)}\Delta t)\right) - \kappa(\theta_{\text{ctrl},i}^{(t+\Delta t)} - \theta_{\text{ctrl},i+1}^{(t+\Delta t)}) + \tau_{\text{joint},i}^{(t+\Delta t)} \\
= &\ (c + \kappa\Delta t)\omega_{i-1}^{(t+\Delta t)} - (\beta + 2c + 2\kappa\Delta t)\omega_i^{(t+\Delta t)} + (c + \kappa\Delta t)\omega_{i+1}^{(t+\Delta t)} \\
&+\kappa(s_{i-1}^{(t)} - 2s_i^{(t)} + s_{i+1}^{(t)}) - \kappa(\theta_{\text{ctrl},i}^{(t+\Delta t)} - \theta_{\text{ctrl},i+1}^{(t+\Delta t)}) + \tau_{\text{joint},i}^{(t+\Delta t)}
\end{aligned}$$

The above formula is impossible to integrate because $\theta_{\text{ctrl},i}^{(t+\Delta t)}$ , $\theta_{\text{ctrl},i+1}^{(t+\Delta t)}$ , $\tau_{\text{joint},i}^{(t+\Delta t)}$ are unknown at time t.

$$\frac{\left|-\kappa\left((\theta_{\text{ctrl},i}^{(t+\Delta t)} - \theta_{\text{ctrl},i}^{(t)}) - (\theta_{\text{ctrl},i+1}^{(t+\Delta t)} - \theta_{\text{ctrl},i+1}^{(t)})\right) + (\tau_{\text{joint},i}^{(t+\Delta t)} - \tau_{\text{joint},i}^{(t)})\right|}{\left|(c + \kappa\Delta t)\omega_{i-1}^{(t+\Delta t)} - (\beta + 2c + 2\kappa\Delta t)\omega_i^{(t+\Delta t)} + (c + \kappa\Delta t)\omega_{i+1}^{(t+\Delta t)} + \kappa(s_{i-1}^{(t)} - 2s_i^{(t)} + s_{i+1}^{(t)}) - \kappa\left(\theta_{\text{ctrl},i}^{(t+\Delta t)} - \theta_{\text{ctrl},i+1}^{(t+\Delta t)}\right) + \tau_{\text{joint},i}^{(t+\Delta t)}\right|} \simeq 0$$

If the above formula is assumed to be true, the following approximation can be used.

$$\begin{aligned}
I\alpha_i^{(t+\Delta t)} = &\ (c + \kappa\Delta t)\omega_{i-1}^{(t+\Delta t)} - (\beta + 2c + 2\kappa\Delta t)\omega_i^{(t+\Delta t)} + (c + \kappa\Delta t)\omega_{i+1}^{(t+\Delta t)} \\
&+\kappa\left(s_{i-1}^{(t)} - 2s_i^{(t)} + s_{i+1}^{(t)}\right) - \kappa\left(\theta_{\text{ctrl},i}^{(t+\Delta t)} - \theta_{\text{ctrl},i+1}^{(t+\Delta t)}\right) + \tau_{\text{joint},i}^{(t+\Delta t)} \\
\simeq &\ (c + \kappa\Delta t)\omega_{i-1}^{(t+\Delta t)} - (\beta + 2c + 2\kappa\Delta t)\omega_i^{(t+\Delta t)} + (c + \kappa\Delta t)\omega_{i+1}^{(t+\Delta t)} \\
&+\kappa\left(s_{i-1}^{(t)} - 2s_i^{(t)} + s_{i+1}^{(t)}\right) - \kappa\left(\theta_{\text{ctrl},i}^{(t)} - \theta_{\text{ctrl},i+1}^{(t)}\right) + \tau_{\text{joint},i}^{(t)} \\
= &\ (c + \kappa\Delta t)\omega_{i-1}^{(t+\Delta t)} - (\beta + 2c + 2\kappa\Delta t)\omega_i^{(t+\Delta t)} + (c + \kappa\Delta t)\omega_{i+1}^{(t+\Delta t)} + \tau_{\kappa,i}^{(t)} + \tau_{\text{joint},i}^{(t)} \\
\omega_i^{(t+\Delta t)} = &\ \omega_i^{(t)} + \alpha_i^{(t+\Delta t)}\Delta t \\
= &\ \omega_i^{(t)} + \frac{\Delta t}{I}\left((c + \kappa\Delta t)\omega_{i-1}^{(t+\Delta t)} - (\beta + 2c + 2\kappa\Delta t)\omega_i^{(t+\Delta t)} + (c + \kappa\Delta t)\omega_{i+1}^{(t+\Delta t)} + \tau_{\kappa,i}^{(t)} + \tau_{\text{joint},i}^{(t)}\right)
\end{aligned}$$

The above formula can be expressed as a matrix formula as follows.

$$\begin{bmatrix} \vdots \\ \omega_{i-1}^{(t+\Delta t)} \\ \omega_{i}^{(t+\Delta t)} \\ \omega_{i+1}^{(t+\Delta t)} \\ \vdots \end{bmatrix} \simeq \begin{bmatrix} \vdots \\ \omega_{i-1}^{(t)} \\ \omega_{i}^{(t)} \\ \omega_{i+1}^{(t)} \\ \vdots \end{bmatrix} + \frac{\Delta t}{I} \begin{bmatrix} \ddots & \vdots & \vdots & \vdots & \ddots \\ \cdots & (c+\kappa\Delta t) & -(\beta+2c+2\kappa\Delta t) & (c+\kappa\Delta t) & \cdots \\ \ddots & \vdots & \vdots & \vdots & \ddots \end{bmatrix} \begin{bmatrix} \vdots \\ \omega_{i-1}^{(t+\Delta t)} \\ \omega_{i}^{(t+\Delta t)} \\ \omega_{i+1}^{(t+\Delta t)} \\ \vdots \end{bmatrix} + \frac{\Delta t}{I} \begin{bmatrix} \vdots \\ \tau_{\kappa,i-1}^{(t)} + \tau_{\text{joint},i-1}^{(t)} \\ \tau_{\kappa,i}^{(t)} + \tau_{\text{joint},i}^{(t)} \\ \tau_{\kappa,i+1}^{(t)} + \tau_{\text{joint},i+1}^{(t)} \\ \vdots \end{bmatrix}$$

In the above vector matrix formula, let us represent the vectors and matrix by the following symbols.

$$\vec{\omega}^{(t)} \equiv \begin{bmatrix} \vdots \\ \omega_{i-1}^{(t)} \\ \omega_{i}^{(t)} \\ \omega_{i+1}^{(t)} \\ \vdots \end{bmatrix}$$

$$\mathbf{P}_{n\times n} \equiv \begin{bmatrix} \ddots & \vdots & \vdots & \vdots & \ddots \\ \cdots & (c+\kappa\Delta t) & -(\beta+2c+2\kappa\Delta t) & (c+\kappa\Delta t) & \cdots \\ \ddots & \vdots & \vdots & \vdots & \ddots \end{bmatrix}$$

$$\vec{\tau}_{\text{rem}}^{(t)} \equiv \begin{bmatrix} \vdots \\ \tau_{\kappa,i-1}^{(t)} + \tau_{\text{joint},i-1}^{(t)} \\ \tau_{\kappa,i}^{(t)} + \tau_{\text{joint},i}^{(t)} \\ \tau_{\kappa,i+1}^{(t)} + \tau_{\text{joint},i+1}^{(t)} \\ \vdots \end{bmatrix}$$

Then, the matrix formula is expressed as follows.

$$\vec{\omega}^{(t+\Delta t)} \simeq \vec{\omega}^{(t)} + \frac{\Delta t}{I}\mathbf{P}_{n\times n}\vec{\omega}^{(t+\Delta t)} + \frac{\Delta t}{I}\vec{\tau}_{\text{rem}}^{(t)}$$

Now, the following approximation can be obtained where $\mathbf{I}_{n\times n}$ is a unit matrix of size $n \times n$.

$$\therefore \vec{\omega}^{(t+\Delta t)} \simeq \left(\mathbf{I}_{n\times n} - \frac{\Delta t}{I}\mathbf{P}_{n\times n}\right)^{-1} \left(\vec{\omega}^{(t)} + \frac{\Delta t}{I}\vec{\tau}_{\text{rem}}^{(t)}\right)$$

#### Correction formula for the rotational inertia of the entire worm

For a floor surface with low friction like water, when numerically integrating the rotational motion of the worm, if $\Delta t > 1 \times 10^{-6}$ sec, the calculation error accumulated for the rotational inertia of the whole worm significantly influenced the calculation result $\omega_i$ and $s_i$. To prevent this, the error is corrected as follows. If $\bar{x}_i \equiv x_i - x_\mathrm{c}$, $\bar{y}_i \equiv y_i - y_\mathrm{c}$, the moment of inertia of the entire worm at time t is as follows by the parallel axis theorem.

$$I_\mathrm{body}^{(t)} = m \sum_i \left( \left( \bar{x}_i^{(t)} \right)^2 + \left( \bar{y}_i^{(t)} \right)^2 \right) + nI$$

If i-rod is approximated as a point particle, the torque applied to the entire worm at time t is as follows where subscription x, y indicates x, y components of the vector. (See numerical integration for translational motion in Supplementary Information)

$$\boldsymbol{\tau}_\mathrm{body}^{(t+\Delta t)} \simeq \frac{\sum_i \left( \bar{x}_i^{(t)} F_{b,i,\mathrm{y}}^{(t)} - \bar{y}_i^{(t)} F_{b,i,\mathrm{x}}^{(t)} \right)}{1 + \frac{b_\perp \Delta t}{M}}$$

If i-rod is approximated as a point particle, the rotational inertia of the whole worm at time t is as follows.

$$L_\mathrm{body}^{(t)} \simeq m \sum_i \left( \bar{x}_i^{(t)} v_{i,\mathrm{y}}^{(t)} - \bar{y}_i^{(t)} v_{i,\mathrm{x}}^{(t)} \right)$$

The predicted value of $\omega_i$ at $t + \Delta t$, $\omega_i^\mathrm{p}$, is calculated by the semi-implicit Euler method (See "Numerical integration of the rotational motion" in Supplementary Information). ($\vec{\omega}^\mathrm{p} = [\cdots \quad \omega_i^\mathrm{p} \quad \cdots]^\mathrm{T}$)

$$\vec{\omega}^\mathrm{p} \simeq \left( \mathbf{I}_{n\times n} - \frac{\Delta t}{I} \mathbf{P}_{n\times n} \right)^{-1} \left( \vec{\omega}^{(t)} + \frac{\Delta t}{I} \vec{\tau}_\mathrm{rem}^{(t)} \right)$$

The predicted value of $s_i$ at $t + \Delta t$ is as follows.

$$s_i^\mathrm{p} = s_i^{(t)} + \omega_i^\mathrm{p} \Delta t$$

Predicted values $x_i^\mathrm{p}$, $y_i^\mathrm{p}$, $\mathbf{v}_i^\mathrm{p}$ for $x_i^{(t+\Delta t)}$, $y_i^{(t+\Delta t)}$, $\mathbf{v}_i^{(t+\Delta t)}$ are calculated from $\mathbf{d}_\mathrm{c}$, $\mathbf{v}_\mathrm{c}$, $s_i^\mathrm{p}$, $\omega_i^\mathrm{p}$ (See "Minimum information required to describe the motion of each rod" in Supplementary Information).

Using $x_i^\mathrm{p}$, $y_i^\mathrm{p}$, $\mathbf{v}_i^\mathrm{p}$, the moment of inertia $I_\mathrm{body}^\mathrm{p}$ and rotational inertia $L_\mathrm{body}^\mathrm{p}$ of the entire worm at time $t + \Delta t$ are calculated as follows. (where $\bar{x}_i^\mathrm{p} \equiv x_i^\mathrm{p} - x_\mathrm{c}^\mathrm{p}$, $\bar{y}_i^\mathrm{p} \equiv y_i^\mathrm{p} - y_\mathrm{c}^\mathrm{p}$, $\mathbf{v}_i = [v_{i,\mathrm{x}} \quad v_{i,\mathrm{y}}]^\mathrm{T}$)

$$I_\mathrm{body}^\mathrm{p} = m \sum_i \left( \left( \bar{x}_i^\mathrm{p} \right)^2 + \left( \bar{y}_i^\mathrm{p} \right)^2 \right) + nI$$

$$L_{\text{body}}^{\text{p}} = m \sum_{i} \left( \bar{x}_i^{\text{p}} v_{i,\text{y}}^{\text{p}} - \bar{y}_i^{\text{p}} v_{i,\text{x}}^{\text{p}} \right)$$

$\omega_i^{(t+\Delta t)}$ is calculated as follows.

$$\omega_i^{(t+\Delta t)} = \omega_i^{\text{p}} + \frac{-\left(L_{\text{body}}^{\text{p}} - L_{\text{body}}^{(t)}\right) + \tau_{\text{body}}^{(t+\Delta t)} \Delta t}{\frac{I_{\text{body}}^{\text{p}} + I_{\text{body}}^{(t)}}{2}}$$

$s_i^{(t+\Delta t)}$ is calculated as follows.

$$s_i^{(t+\Delta t)} = s_i^{(t)} + \omega_i^{(t+\Delta t)} \Delta t$$

By correcting the rotational inertia for the whole worm, numerical integration of the rotational motion of the worm was well calculated even for cases when $\Delta t > 1 \times 10^{-6}$ sec, as if $\Delta t \leq 1 \times 10^{-6}$ sec.

#### Locomotion propulsion mechanism

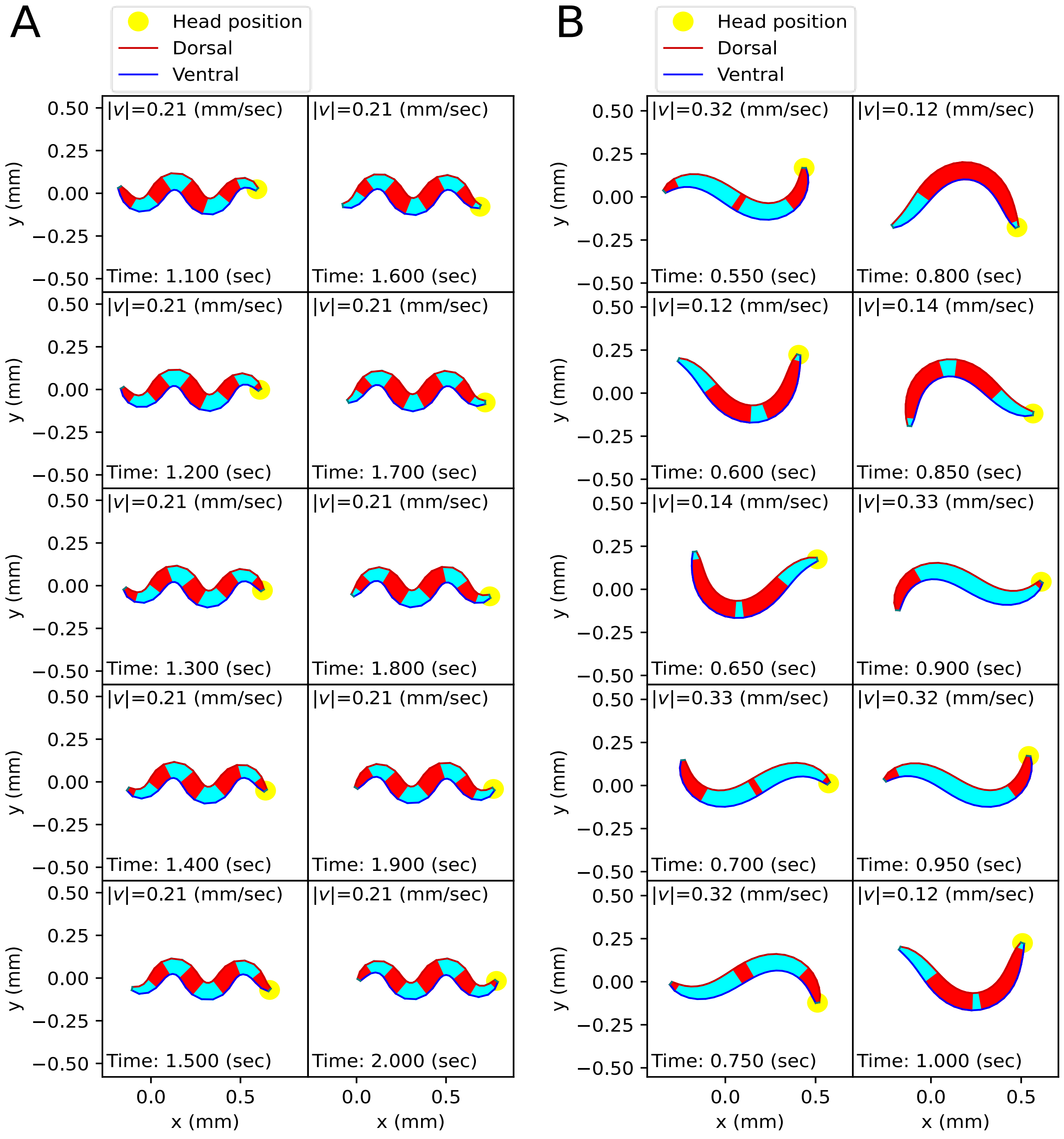


**Figure S3**. Locomotion propulsion mechanism. During the worm's locomotion, the parts receiving force in the direction of progress are marked in red, and the parts receiving force in the opposite direction are marked in cyan. (A) Crawling propulsion mechanism. During crawling, the worm gains propulsion from the body parts diagonal to the direction of progress. (B) Swimming propulsion mechanism. During swimming, the worm generates propulsion when it bends into a C-shape. While the worm's speed is almost constant during crawling, it is not constant during swimming.

**Proper Selection of Friction Coefficients**

When using the vertical and horizontal friction coefficients $b_{\text{agar},\perp}$, $b_{\text{agar},\parallel}$ on agar, as proposed in the previous work (Boyle et al., 2012), the trajectory of the escaping behavior was not accurately replicated. Therefore, we sought appropriate friction coefficients necessary for replicating the escaping behavior. We calculated new vertical and horizontal friction coefficients $b_{\eta,\perp} = \eta b_{\text{agar},\perp}$, $b_{\eta,\parallel} = \eta b_{\text{agar},\parallel}$ by multiplying scaling factor $\eta$ to the agar friction coefficients $b_{\text{agar},\perp}$, $b_{\text{agar},\parallel}$ of the previous work (Boyle et al., 2012). Let us denote the set of a quantity for all pairs $(i,t)$ of index $i$ and time $t$ as $\{*\}_{i,t}$. When the kymogram input $\{\theta_{\text{ctrl},i}^{(t)}\}_{i,t}$ was same as Fig. 3A, we observed how the trajectory of the escaping behavior changes with $\eta$ (Fig. S4A) and analyzed representative values for each trajectory as follows (Fig. S4B) (Note that when $\eta$ was less than or equal to $10^{-6}$, the time-step ($\Delta t$) was set to $10^{-6}$ sec for higher accuracy of simulation). Let us denote the average of a quantity for all pairs $(i,t)$ as $\langle * \rangle_{i,t}$. When $\eta$ was $1 \sim 10^{-9}$, the smaller $\eta$, the smaller $E_\theta = \left\langle \left| \theta_i^{(t)} - \theta_{\text{ctrl},i}^{(t)} \right| \right\rangle_{i,t}$ was. The reduction in $E_\theta$ when $\eta$ changed from $10^{-2}$ to $10^{-9}$ was about 3% of the reduction when $\eta$ changed from 1 to $10^{-9}$. When $\eta$ was between 1 and $10^{-9}$, even if $\eta$ decreased, $E_\theta$ did not fall below 0.125 (rad). This is because there was a time delay between the input $\theta_{\text{ctrl},i}^{(t)}$ and the response $\theta_i^{(t)}$. As $\eta$ decreased from 1 to $10^{-2}$, the total traveled distance of the worm ($\sum_t \left| \mathbf{v}_\text{c}^{(t)} \Delta t \right|$) and the total absolute angle change ($\mathcal{S} = \sum_{t=0}^{T-\Delta t} \left| \left\langle s_i^{(t+\Delta t)} \right\rangle_i - \left\langle s_i^{(t)} \right\rangle_i \right|$) increased, and the trajectory became more similar to the experimental video (Broekmans et al., 2016). When $\eta$ was between $10^{-2}$ and $10^{-6}$, the worm's trajectory was almost identical, and thus the total traveled distance, $\mathcal{S}$, and the pattern of $\left| \mathbf{F}_{b,i}^{(t)} \right|$ (Fig. S5) were similar across trajectories. If $\eta$ was smaller than $10^{-6}$, the worm's total traveled distance decreased and $\mathcal{S}$ increased, and the trajectory was no longer similar to the experimental video. This was because a too small friction coefficients hindered the worm from obtaining enough propulsive force from the ground (Fig. S5).

The same method with the kymogram input $\{\theta_{\text{ctrl},i}^{(t)}\}_{i,t}$ same as Fig. 3B used to analyze the effect of $\eta$ on the trajectory of the escaping behavior was applied to analyze the trajectory of the delta-turn according to $\eta$ (Fig. S6A). When $\eta$ was $1 \sim 10^{-9}$, the smaller $\eta$, the smaller $E_\theta$ was (Fig. S6B). The reduction in $E_\theta$ when $\eta$ changed from $10^{-2}$ to $10^{-9}$ was about 3% of the reduction when $\eta$ changed from 1 to $10^{-9}$. When $\eta$ was between 1 and $10^{-9}$, even if $\eta$ decreased, $E_\theta$ did not fall below 0.15

(rad). As $\eta$ decreased from 1 to $10^{-2}$, $\mathcal{S}$ increased, and the trajectory became more similar to the experimental video. When $\eta$ was between $10^{-2}$ and $10^{-6}$, the worm's trajectory was almost identical, and thus the total traveled distance, $\mathcal{S}$, and the pattern of $\left|\mathbf{F}_{b,i}^{(t)}\right|$ (Fig. S7) were similar across trajectories. If $\eta$ was smaller than $10^{-6}$, the worm's total traveled distance decreased and $\mathcal{S}$ increased, and the trajectory was no longer similar to the experimental video. This is because a too small friction coefficients hindered the worm from obtaining enough propulsive force from the ground (Fig. S7).

In conclusion, when the ratio between vertical and horizontal friction coefficients was constant at $40 (= b_{\eta,\perp}/b_{\eta,\parallel} = b_{\text{agar},\perp}/b_{\text{agar},\parallel})$, selecting an appropriate $\eta$ value (between $10^{-2}$ and $10^{-6}$) was crucial for replicating the trajectory of sequenced locomotive behavior. Therefore, we chose $b_{\eta,\perp}$, $b_{\eta,\parallel}$ ($\eta = 10^{-2}$) as the friction coefficients that sufficiently reduce $E_\theta$ among those closest to the agar friction coefficients of the previous work (Boyle et al., 2012) for both escaping behavior and delta-turn.

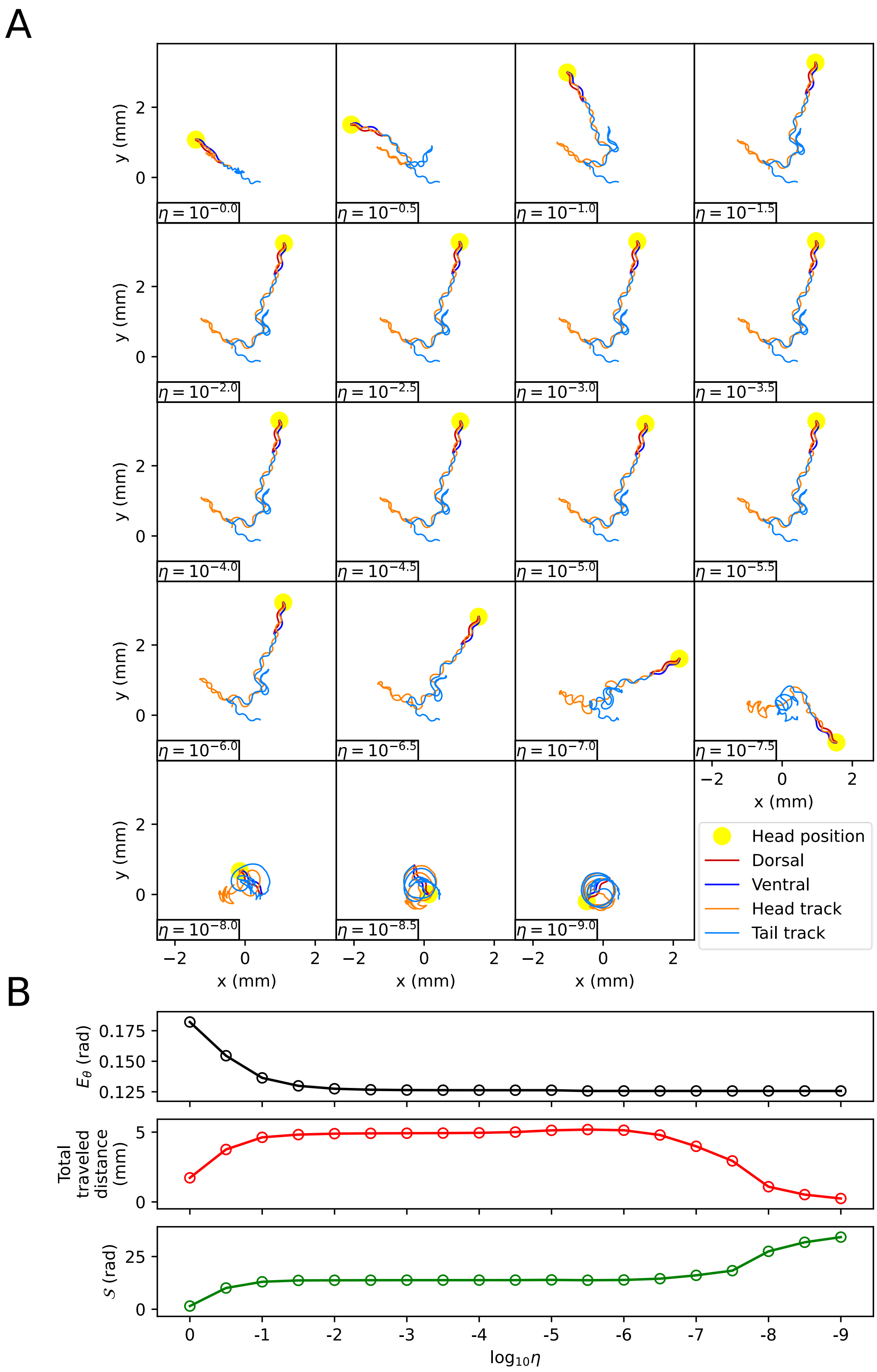
A
B
Head position
Dorsal
Ventral
Head track
Tail track
x (mm)
y (mm)
$\eta = 10^{-0.0}$
$\eta = 10^{-0.5}$
$\eta = 10^{-1.0}$
$\eta = 10^{-1.5}$
$\eta = 10^{-2.0}$
$\eta = 10^{-2.5}$
$\eta = 10^{-3.0}$
$\eta = 10^{-3.5}$
$\eta = 10^{-4.0}$
$\eta = 10^{-4.5}$
$\eta = 10^{-5.0}$
$\eta = 10^{-5.5}$
$\eta = 10^{-6.0}$
$\eta = 10^{-6.5}$
$\eta = 10^{-7.0}$
$\eta = 10^{-7.5}$
$\eta = 10^{-8.0}$
$\eta = 10^{-8.5}$
$\eta = 10^{-9.0}$
$E_\theta$ (rad)
Total traveled distance (mm)
$S$ (rad)
$\log_{10}\eta$

**Figure S4**: The effect of the scaling factor $\eta$ of the friction coefficients on the escaping behavior (A) The trajectory of the worm for each scaling factor $\eta$. (B) Characteristics of the trajectory. The top graph represents $E_\theta = \left\langle \left| \theta_i^{(t)} - \theta_{\mathrm{ctrl},i}^{(t)} \right| \right\rangle_{i,t}$. The middle graph shows the total traveled distance of the worm. The bottom graph represents the total absolute angle change ($\mathcal{S} = \sum_{t=0}^{T-\Delta t} \left| \left\langle s_i^{(t+\Delta t)} \right\rangle_i - \left\langle s_i^{(t)} \right\rangle_i \right|$, where $T$ is the total time of the experimental video).

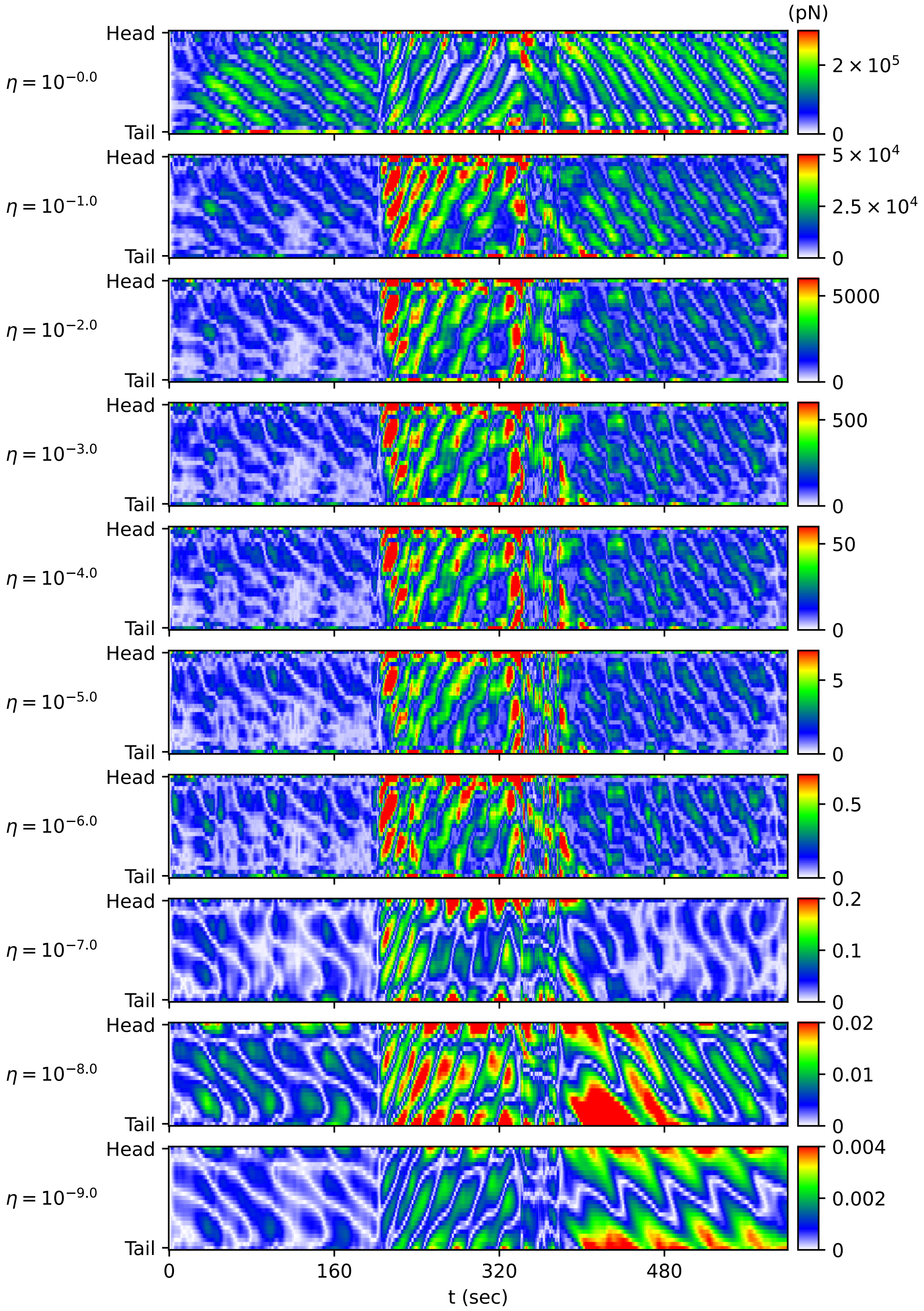


**Figure S5**: The magnitude of the frictional force $\left|\mathbf{F}_{b,i}^{(t)}\right|$ during the escaping behavior depending on the scaling factor $\eta$ of the friction coefficient. The color of each point in the heatmap represents the value of $\left|\mathbf{F}_{b,i}^{(t)}\right|$.

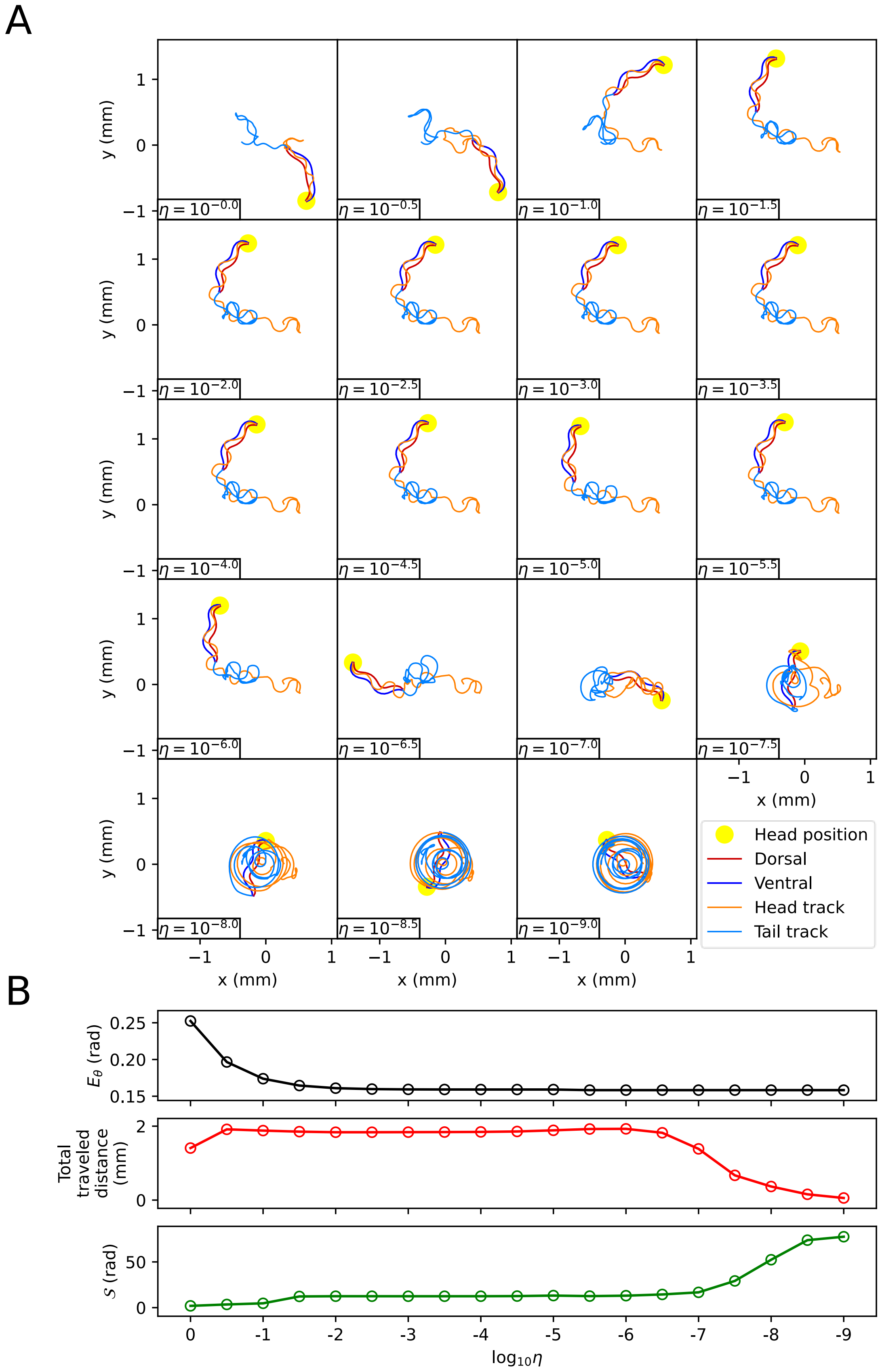
A
B
y (mm)
x (mm)
$\eta = 10^{-0.0}$
$\eta = 10^{-0.5}$
$\eta = 10^{-1.0}$
$\eta = 10^{-1.5}$
$\eta = 10^{-2.0}$
$\eta = 10^{-2.5}$
$\eta = 10^{-3.0}$
$\eta = 10^{-3.5}$
$\eta = 10^{-4.0}$
$\eta = 10^{-4.5}$
$\eta = 10^{-5.0}$
$\eta = 10^{-5.5}$
$\eta = 10^{-6.0}$
$\eta = 10^{-6.5}$
$\eta = 10^{-7.0}$
$\eta = 10^{-7.5}$
$\eta = 10^{-8.0}$
$\eta = 10^{-8.5}$
$\eta = 10^{-9.0}$
Head position
Dorsal
Ventral
Head track
Tail track
$E_\theta$ (rad)
Total traveled distance (mm)
$S$ (rad)
$\log_{10}\eta$

**Figure S6**: The impact of the scaling factor $\eta$ of the friction coefficients on the delta-turn (A) The trajectory of the worm for each scaling factor $\eta$. (B) Characteristics of the trajectory. The top graph represents $E_\theta = \left\langle \left| \theta_i^{(t)} - \theta_{\mathrm{ctrl},i}^{(t)} \right| \right\rangle_{i,t}$. The middle graph shows the total traveled distance of the worm. The bottom graph represents the total absolute angle change ($\mathcal{S} = \sum_{t=0}^{T-\Delta t} \left| \left\langle s_i^{(t+\Delta t)} \right\rangle_i - \left\langle s_i^{(t)} \right\rangle_i \right|$).

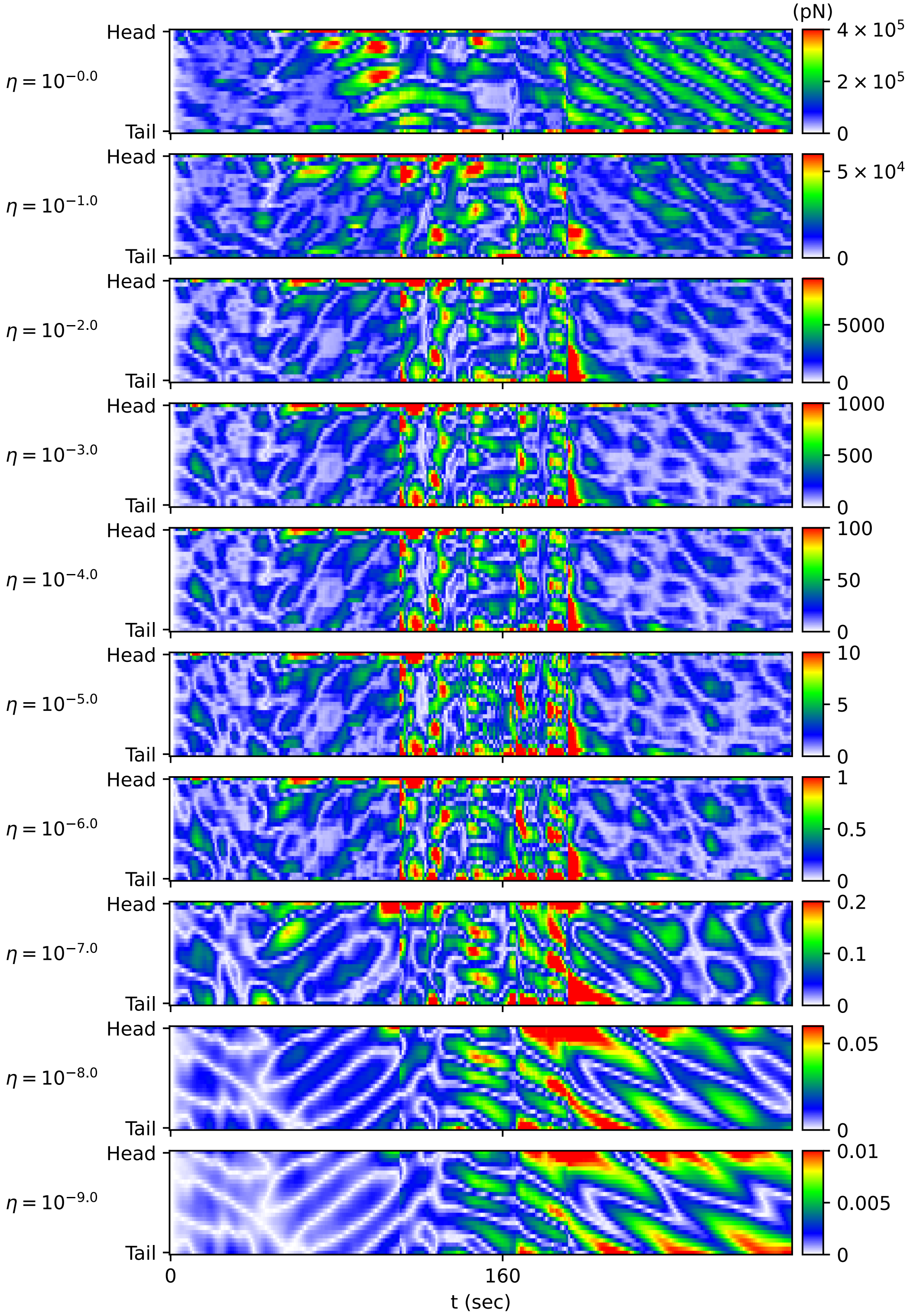


**Figure S7**: The magnitude of the frictional force $\left|\mathbf{F}_{b,i}^{(t)}\right|$ during the delta-turn depending on the scaling factor $\eta$ of the friction coefficient. The color of each point in the heatmap represents the value of $\left|\mathbf{F}_{b,i}^{(t)}\right|$.

#### Transition of body shape from water to agar

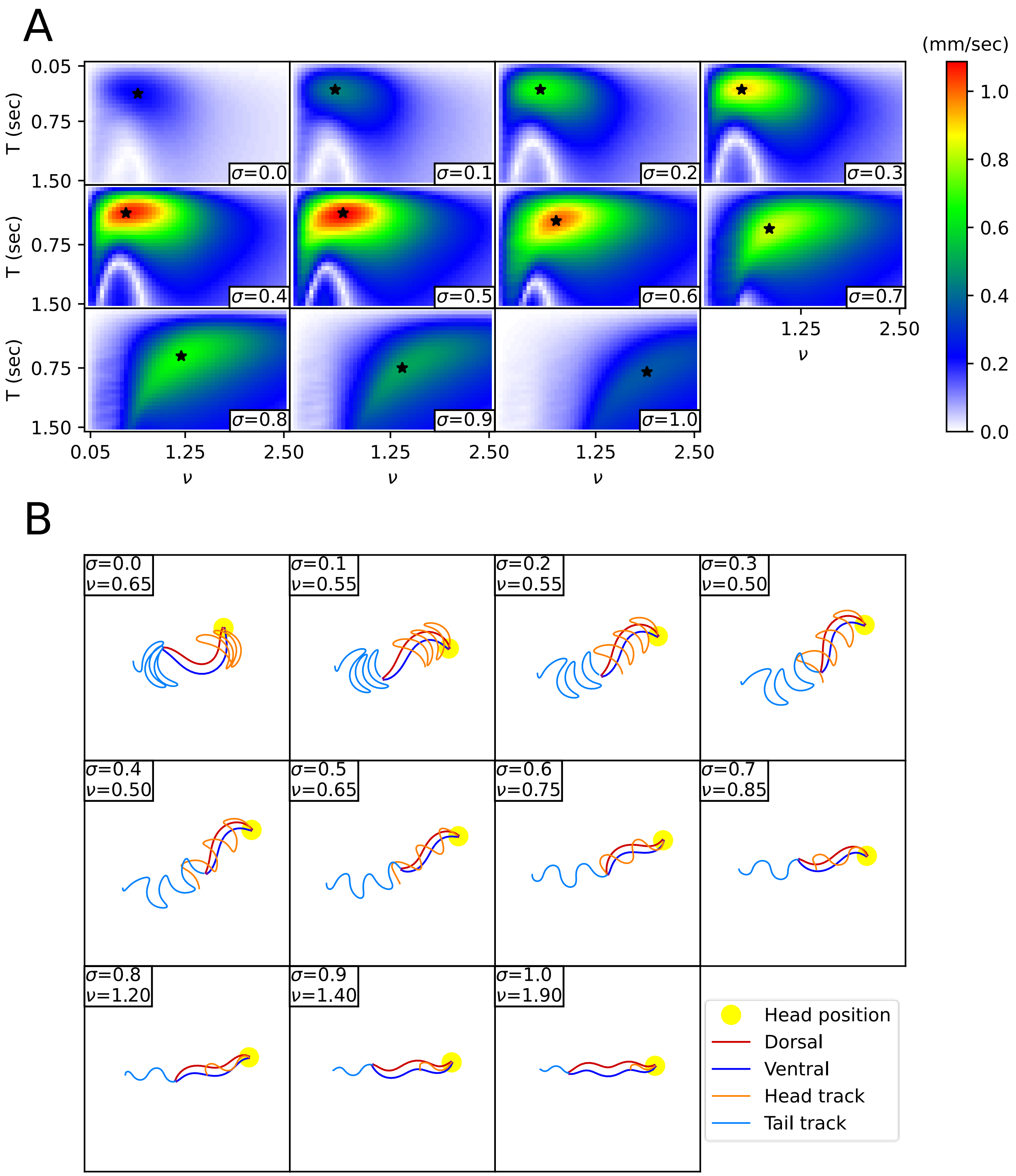


**Figure S8**. Transition of body shape from water ($\sigma = 0$) to agar ($\sigma = 1$). (A) The average velocity of the worm according to the wavenumber $\nu$ and period $T$ to the friction coefficient by $\sigma$. The star marker denotes the pair of $(\nu, T)$ that maximizes the average velocity of the worm (i.e., optimal $(\nu, T)$). As $\sigma$ increases and the friction coefficient gradually increases, the optimal $(\nu, T)$ transitions from the upper left to the lower right. (B) The body shape and locomotion pattern due to the pair of $(\nu, T)$ that maximizes the average velocity of the worm for the friction coefficient by $\sigma$.

#### Effect of environmental index on sequenced locomotive behavior

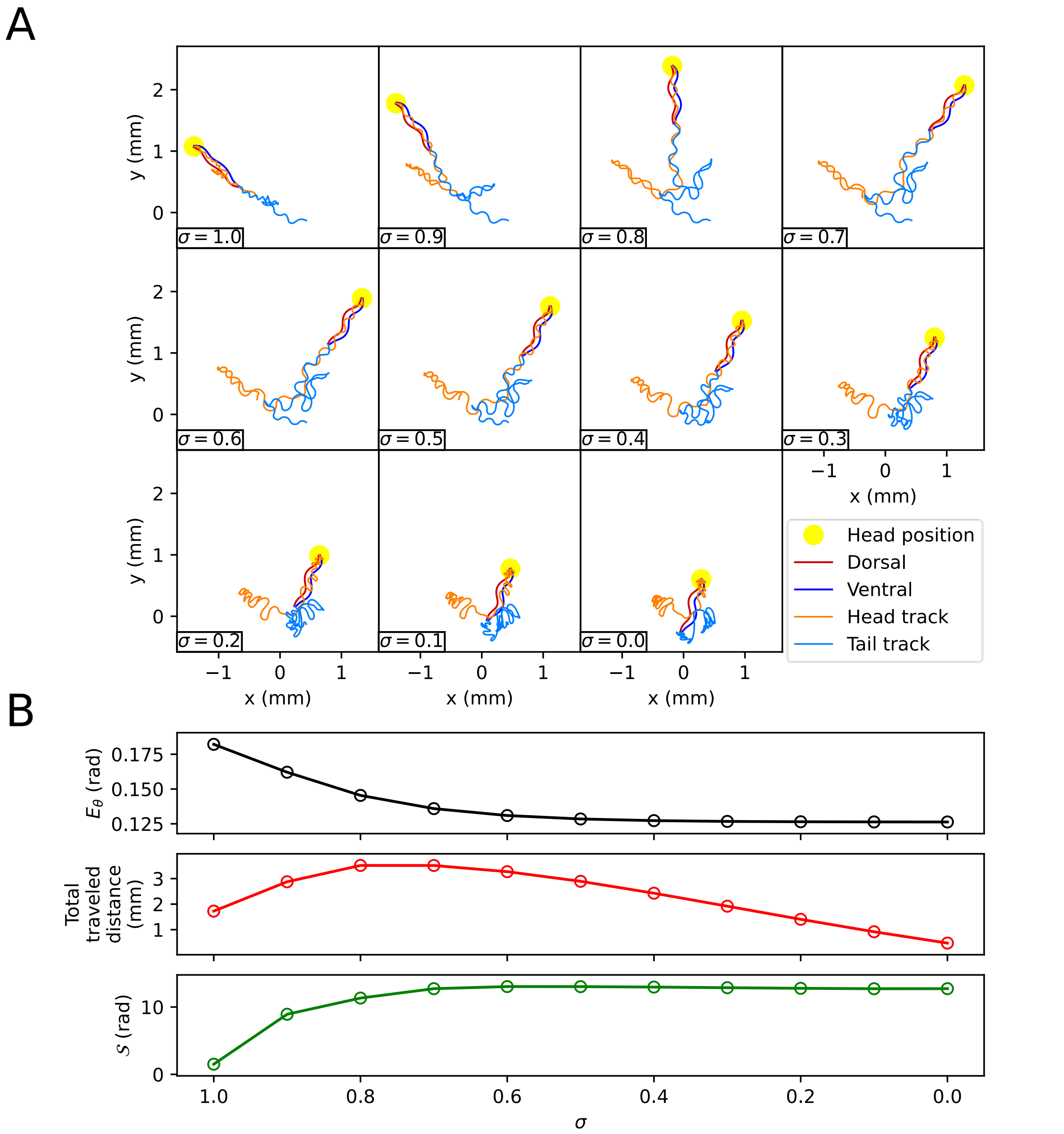


**Figure S9**: The effect of the environmental index $\sigma$ on escaping behavior (A) The trajectory of the worm depending on the environmental index $\sigma$. (B) Characteristics of the trajectory. The top graph represents $E_\theta = \left\langle \left| \theta_i^{(t)} - \theta_{\mathrm{ctrl},i}^{(t)} \right| \right\rangle_{i,t}$. The middle graph shows the total traveled distance of the worm. The bottom graph represents the total absolute angle change ($\mathcal{S} = \sum_{t=0}^{T-\Delta t} \left| \left\langle s_i^{(t+\Delta t)} \right\rangle_i - \left\langle s_i^{(t)} \right\rangle_i \right|$.

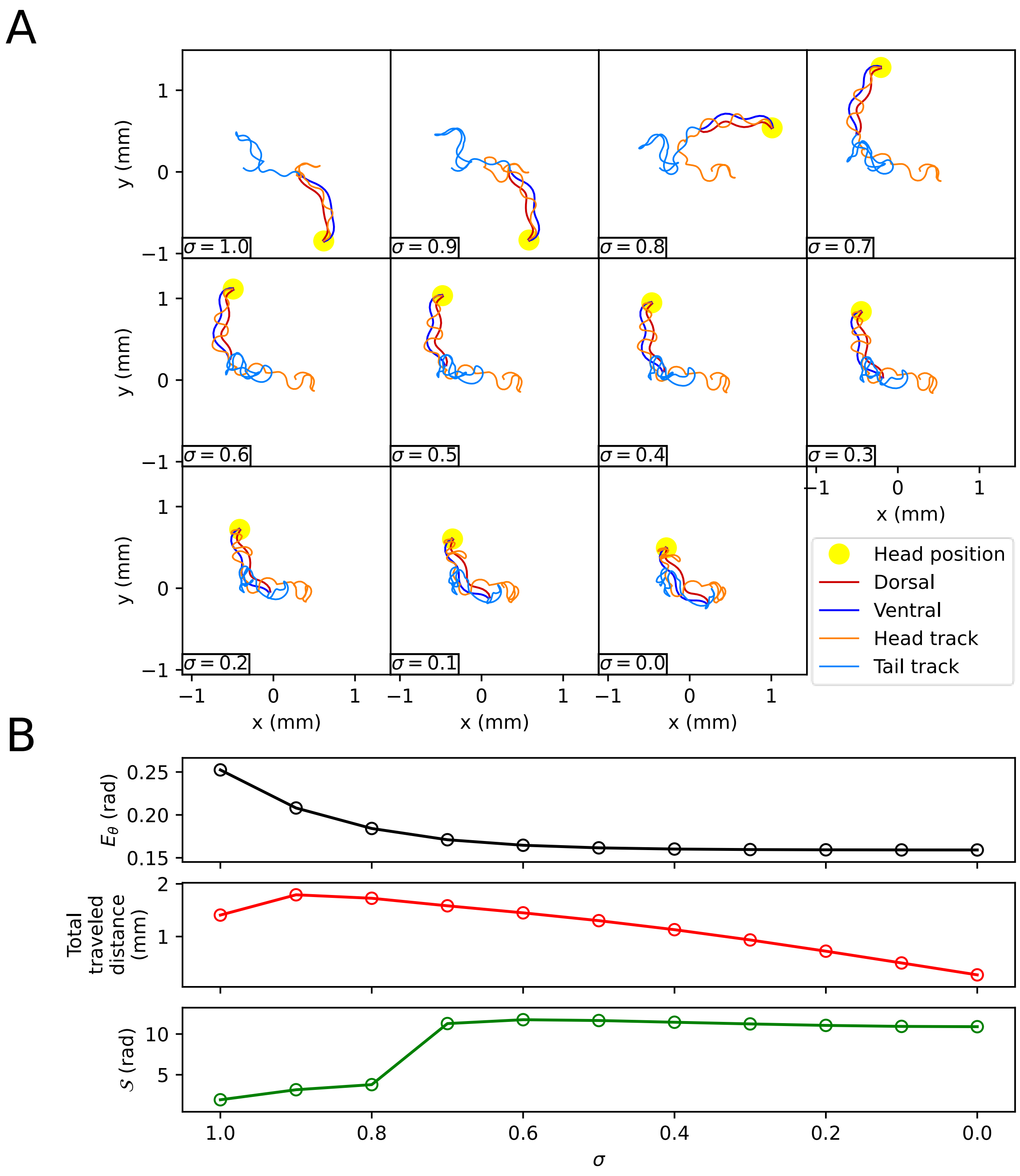


**Figure S10**: The effect of the environmental index $\sigma$ on the delta-turn (A) The trajectory of the worm for each environmental index $\sigma$. (B) Characteristics of the trajectory. The top graph represents $E_\theta = \left\langle \left| \theta_i^{(t)} - \theta_{\mathrm{ctrl},i}^{(t)} \right| \right\rangle_{i,t}$. The middle graph shows the total traveled distance of the worm. The bottom graph represents the total absolute angle change ($\mathcal{S} = \sum_{t=0}^{T-\Delta t} \left| \left\langle s_i^{(t+\Delta t)} \right\rangle_i - \left\langle s_i^{(t)} \right\rangle_i \right|$).

#### Process of Defining Behavioral Categories

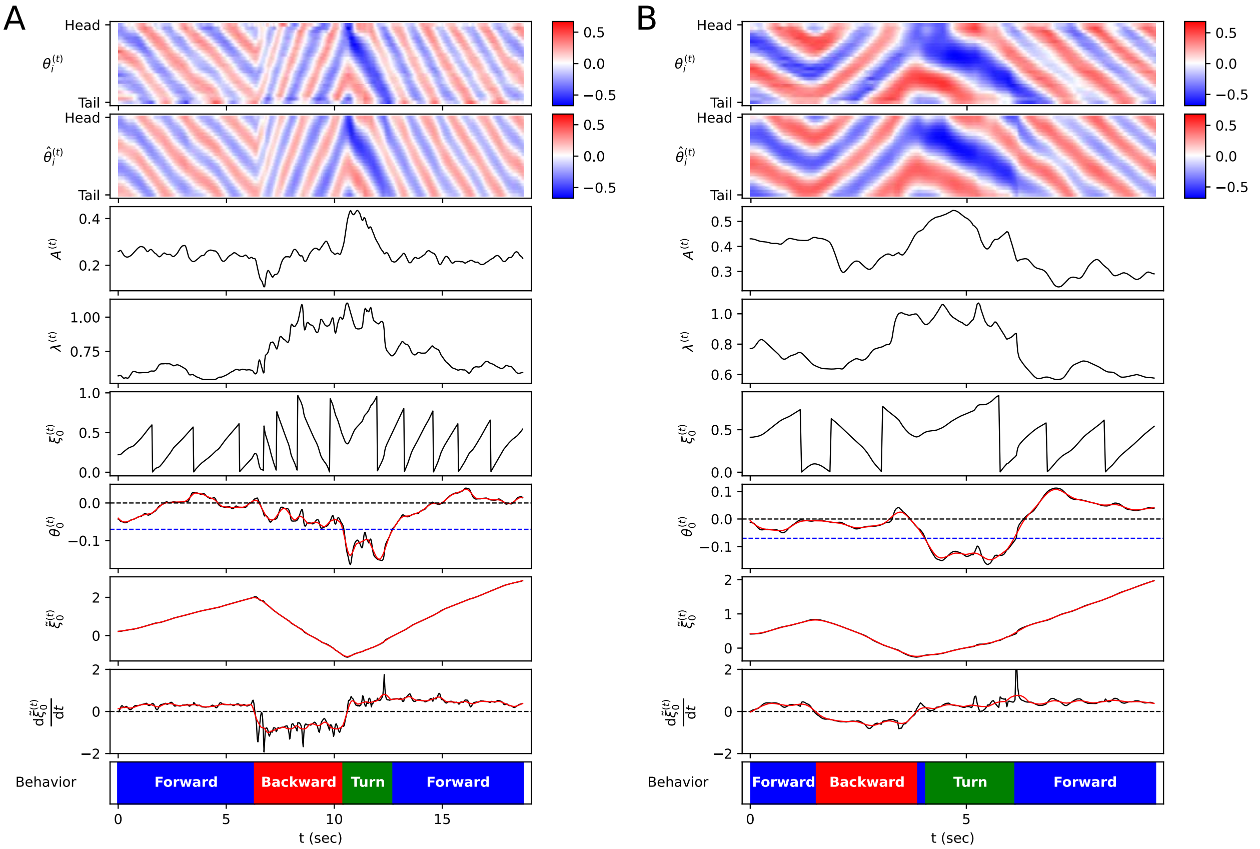


**Figure S11**. Process of defining behavioral categories. (A) Escaping behavior. The top heatmap represents the body angle $\theta_i^{(t)}$. The second panel from the top is the sine fitting of $\theta_i^{(t)}$, denoted as $\hat{\theta}_i^{(t)}$. The third to the eighth panels present the parameters derived from the sine fitting (amplitude $A^{(t)}$, wavelength $\lambda^{(t)}$, phase $\xi_0^{(t)}$, body angle bias $\theta_0^{(t)}$), and values derived from these parameters ($\tilde{\xi}_0^{(t)}$, $\frac{\mathrm{d}\tilde{\xi}_0^{(t)}}{\mathrm{d}t}$). The black solid line represents the actual value, and the red solid line represents the smoothed value. The bottom panel shows the behavior classification calculated from $\theta_0^{(t)}$ and $\frac{\mathrm{d}\tilde{\xi}_0^{(t)}}{\mathrm{d}t}$ (blue: forward locomotion, red: backward locomotion, green: turn). (B) Delta-turn.

#### Supplementary Information References


Boyle, J. H., Berri, S., & Cohen, N. (2012). Gait Modulation in C. elegans: An Integrated Neuromechanical Model. *Frontiers in Computational Neuroscience*, *6*. https://doi.org/10.3389/fncom.2012.00010

Broekmans, O. D., Rodgers, J. B., Ryu, W. S., & Stephens, G. J. (2016). Resolving coiled shapes reveals new reorientation behaviors in C. elegans. *eLife*, *5*, e17227. https://doi.org/10.7554/eLife.17227

Butcher, J. C. (2004). *Numerical Methods for Ordinary Differential Equations*. John Wiley & Sons.

Reina, A., Subramaniam, A. B., Laromaine, A., Samuel, A. D. T., & Whitesides, G. M. (2013). Shifts in the Distribution of Mass Densities Is a Signature of Caloric Restriction in Caenorhabditis elegans. *PLOS ONE*, *8*(7), e69651. https://doi.org/10.1371/journal.pone.0069651


#### Video legends

**Video S1**. Reproduced escaping behavior of experimental video(Broekmans et al., 2016). Left: Original video. Right: Reproduced video by ElegansBot.

**Video S2**. Reproduced delta-turn of experimental video(Broekmans et al., 2016). Left: Original video. Right: Reproduced video by ElegansBot.
